# Cholinergic synaptic plasticity shapes resilience and vulnerability to tau

**DOI:** 10.1101/2025.05.27.656174

**Authors:** Kate M. Onuska, Hayley R.C. Shanks, Lauren A. Devito, Qi Qi, Alycia M. Crooks, Liliana Germán-Castelán, Geoffrey N. Ngo, Roy A.M. Haast, Tallulah S. Andrews, Kayla M. Williams, Flávio H. Beraldo, Ting Qiu, Alfonso Fajardo-Valdez, Jordana Remz, Alfie Wearn, Gary R. Turner, Etienne Aumont, Jonathan D. Thiessen, Matthew S. Fox, Justin W. Hicks, Timothy J. Bussey, Lisa M. Saksida, Jennifer Tremblay-Mercier, John C.S. Breitner, Jean-Paul Soucy, Judes Poirier, Marc-André Bedard, Sylvia Villeneuve, Vania F. Prado, Marco A.M. Prado, R. Nathan Spreng, Taylor W. Schmitz, the PREVENT-AD Research Group.

## Abstract

Synaptic dysfunction is a hallmark of Alzheimer’s disease (AD). Yet due to their plasticity, synapses may also adapt to early AD pathology. Here, we demonstrate that cholinergic neurons mount a presynaptic response to tau pathology in the living human brain. Using multi-tracer positron emission tomography in cognitively normal older adults at risk for AD, we observe that cholinergic neurons increase presynaptic vesicular acetylcholine transporter (VAChT) protein levels when colocalized to tau, but not amyloid. Notably, stronger VAChT responses were associated with cognitive resilience over a decade. Whole-brain single-nucleus RNA sequencing in human and mouse tissue reveal that cholinergic neurons are enriched for a plasticity gene-network anchored to the microtubule-associated protein tau (*MAPT*) gene. In mice, forebrain-specific deletion of VAChT impairs cortical plasticity and hippocampal structural integrity. Overall, our findings identify cholinergic synaptic plasticity, and its failure, as a fundamental mechanism of resilience and vulnerability to tau in presymptomatic AD.

## Main

Synaptic dysfunction is among the first detectable features of Alzheimer’s disease (AD)^1,2^, and the strongest correlate of early cognitive decline^3-7^. Yet synapses are also among the brain’s most dynamic and adaptable structures. Synapses can rebalance their strength and connectivity through plasticity, a form of cellular adaptation that supports learning under normal conditions and helps stabilize circuits under sustained stress^8-11^. Such adaptive capacity may even explain how many individuals maintain cognitive function despite years of accumulating AD pathology^12^.

Whether synapses mount such compensatory responses to specific AD pathologies in the living human brain remains unknown. Prevailing models of AD assume a passive trajectory: β-amyloid (Aβ) and tau pathologies accumulate, circuits fail, and cognition declines. But if synaptic adaptation occurs before neurodegeneration, then it may not only delay decline, but also determine which individuals resist it^13,14^. The challenge is to capture these early synaptic changes *in vivo*, and disentangle whether they reflect compensation to pathophysiological vulnerabilities.

Here we leverage molecular imaging to quantify cell-type-specific presynaptic function alongside regional tau and Aβ pathology in healthy older adults at elevated risk for AD. We show that basal forebrain (BF) cholinergic synapses mount an adaptive response to early tau pathology, but not Aβ. We demonstrate that this response is associated with longitudinal cognitive resilience and the spatial extent of tau pathology in the brain. Supported by transcriptomic and multimodal neuroimaging work across species, our results position cholinergic synaptic plasticity as a homeostatic buffer to tau, and its failure as a tipping point toward neurodegeneration.

### Imaging cholinergic presynaptic function *in vivo*

Extensive neuropathological staging and neuroimaging work points to the cholinergic BF as one of the first sites of pathology and neurodegeneration in the progression of AD^15-18^. More recent studies have demonstrated that longitudinal BF atrophy in at-risk older adults predicts the progression of cortical neurodegeneration^19^, which reflects the spatial profile of cholinergic presynaptic loss observed in AD patients^20,21^. Parallel lines of evidence suggest that BF cholinergic synapses are essential for promoting Hebbian plasticity^22-27^ and stabilizing cellular homeostasis^28-32^ in adult neural circuits. Together, these findings suggest that the BF cholinergic system’s selective vulnerability may partly reflect early presynaptic adaptations to sustain plasticity and network function under increasing pathological load.

To explore BF cholinergic synaptic plasticity in healthy older adults at elevated risk for AD, we collected [^18^F]-fluoroethoxy-benzovesamicol ([^18^F]-FEOBV) positron emission tomography (PET) data in 64 individuals enrolled in the PRe-symptomatic EValuation of Experimental or Novel Treatments for Alzheimer’s Disease (PREVENT-AD) program (**Table 1**, main cohort), an ongoing, longitudinal observational study of cognitively unimpaired individuals recruited on the basis of a first-degree family history of AD^33,34^ (*Methods*). [^18^F]-FEOBV is one of a limited number of PET tracers that permits direct *in vivo* imaging of synapses in humans^35^. It is a highly specific, high-affinity radioligand^36-38^ for the vesicular acetylcholine transporter (VAChT), a protein exclusively produced by cholinergic neurons and most abundantly expressed within the presynaptic nerve terminals^39^. Because VAChT is responsible for packaging acetylcholine into synaptic vesicles to facilitate cholinergic neurotransmission^40^, [^18^F]-FEOBV uptake provides a cell-type specific measure of cholinergic presynaptic function. To investigate whether adaptations in cholinergic presynaptic function track with early pathological load, we also acquired tau ([^18^F]-AV1451) and β-amyloid (Aβ; [^18^F]-NAV4694) PET data in each main cohort participant (**Table 1**, **Extended Data Fig. 1**, main cohort).

**Table 1.**
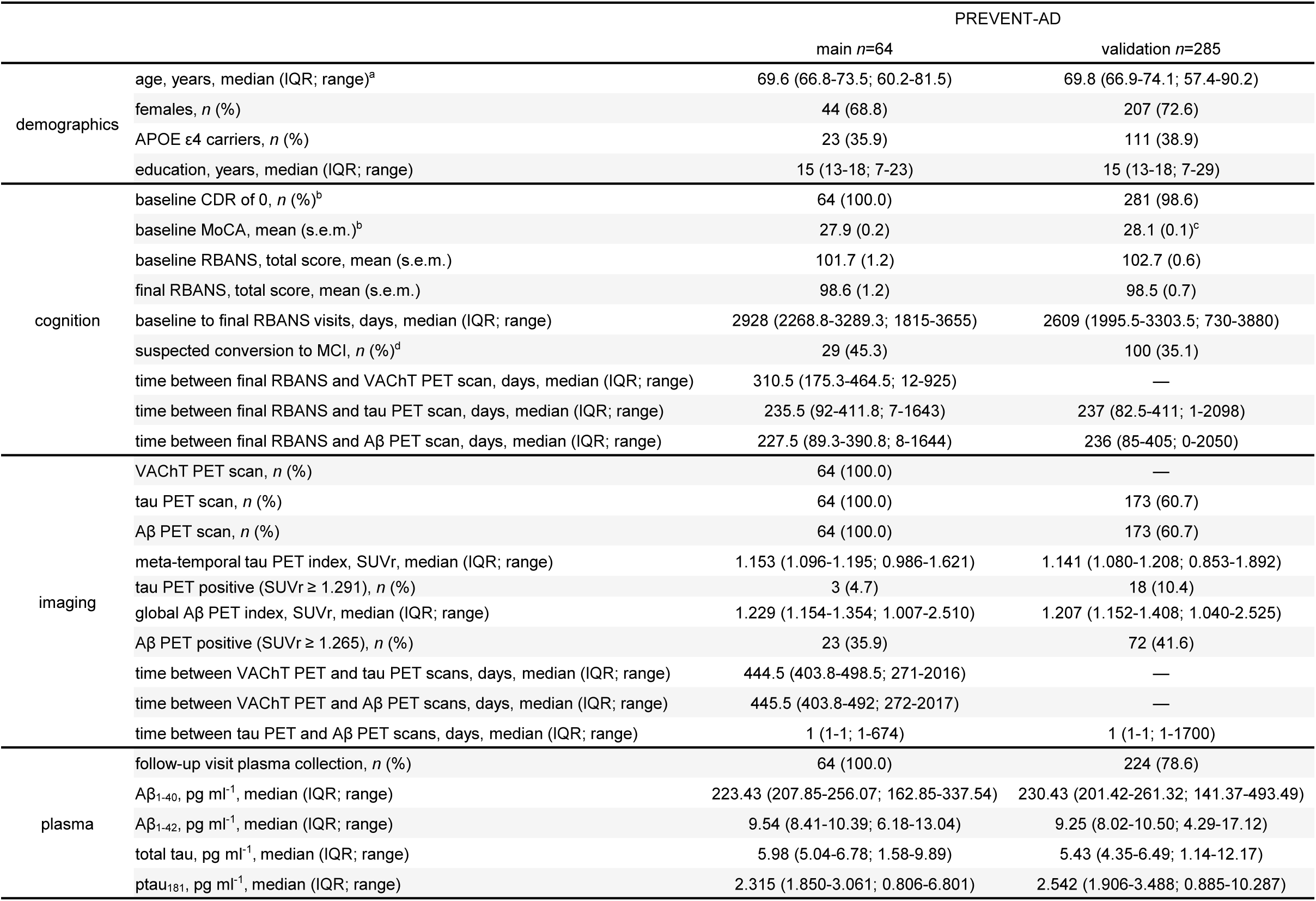
PREVENT-AD main and validation cohort demographic information. ^a^age at final RBANS visit. ^b^information on inclusion criteria for individuals with a baseline CDR > 0 or MoCA < 26 can be found in *Methods*. ^c^missing MoCA data for one participant, mean (standard error of the mean) calculated from *n*=284. ^d^suspected conversion status as of 2025; information on evaluation of clinical progression can be found in *Methods*. Continuous data are presented as the mean (standard error of the mean) or median (interquartile range; range). Categorical data are presented as the number of participants (percentage). Aβ, β-amyloid; APOE ε4, apolipoprotein E ε4 allele; CDR, clinical dementia rating; MCI, mild cognitive impairment; MoCA, Montreal cognitive assessment; ptau_181_, phosphorylated tau at threonine-181; RBANS, repeatable battery for the assessment of neuropsychological status; SUVr, standardized uptake value ratio; VAChT, vesicular acetylcholine transporter.

### Mapping personalized brain profiles of tau and Aβ

We anticipated that the spatial profiles of tau and Aβ pathology, as well as their potential interdependencies with VAChT, would likely exhibit considerable variation across healthy older adults at risk for AD^41-44^. We therefore adapted a normative deviation mapping approach^45,46^ to capture each participant’s unique pattern of tau and Aβ pathology in the brain (*Methods*). Using an iterative, leave-one-out strategy, we generated voxel-wise population median (*M-i*) and scaled median absolute deviation (*MAD-i*) reference images across the main cohort, ensuring statistical independence between each participant’s PET images and their corresponding reference images. Supporting the robustness of this strategy, we observed consistently high correlations across all pairwise combinations of *M-i* or *MAD-i* reference images (**Supplemental Table 1**). For each participant, we identified voxels with tau-PET or Aβ-PET uptake values falling above or below the 95% confidence interval (CI) of their respective *M-i* and *MAD-i* reference voxels. These voxels were classified as regions of high (i.e., >U95) or low (i.e., <L95) pathology, while voxels within the 95% CIs were assigned to intermediate load regions (i.e., L95–M and M–U95) (**Fig. 1a**). Overall, our normative deviation mapping approach identifies individualized profiles of tau and Aβ which are significantly higher or lower than expected in a population of healthy older adults at elevated risk for AD (**Extended Data Fig. 2,3**).

**Fig. 1.**
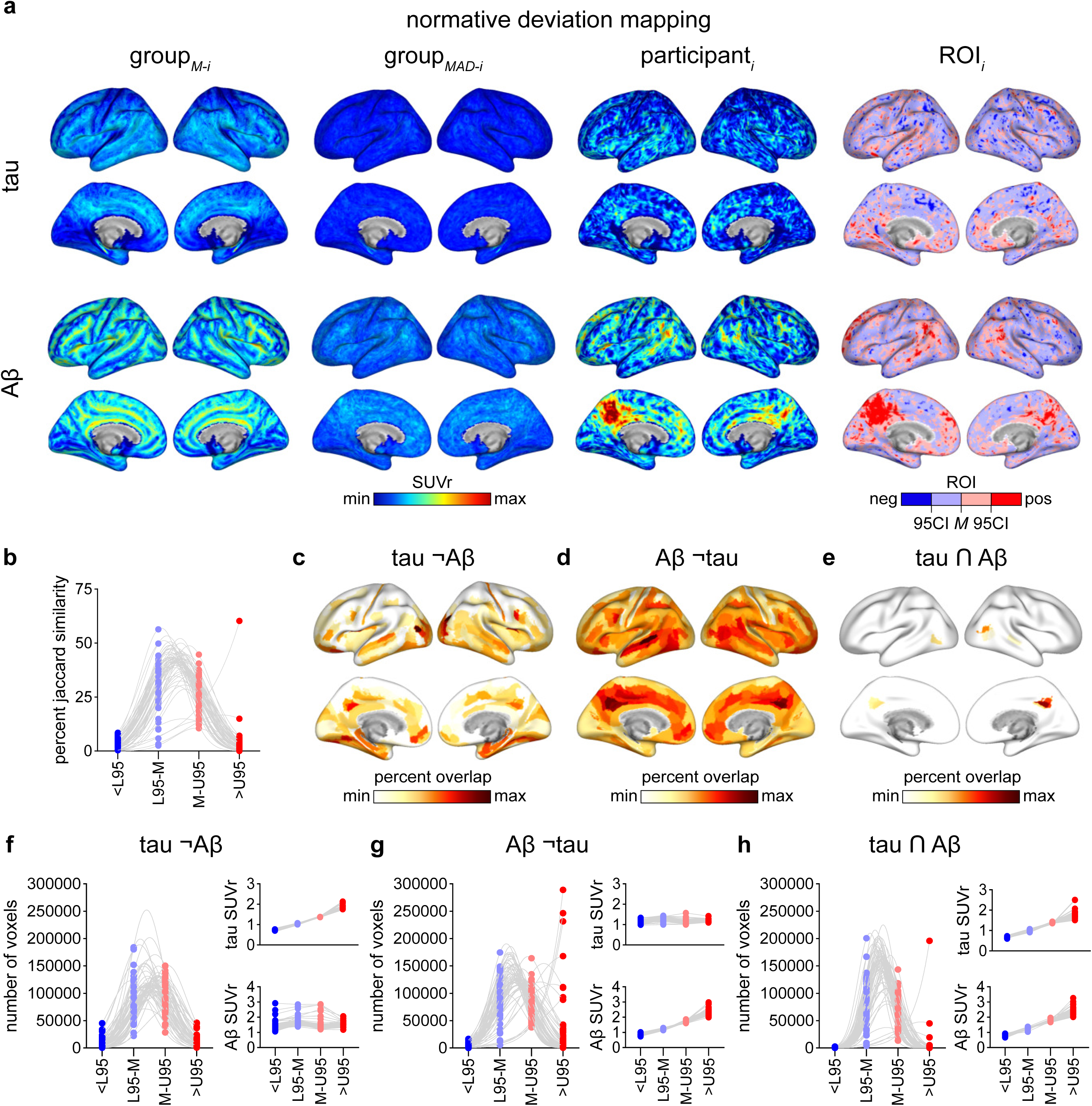
Individualized mapping of tau and Aβ in healthy older adults at elevated risk for Alzheimer’s disease. **a**, Representative normative median deviation mapping steps for standardized uptake value ratio (SUVr) tau-PET (*top*) and Aβ-PET (*bottom*) images. From left to right: (1) population median reference images (tau*_M_*_-_*_i_*: min-max=0.9–2.4; Aβ*_M_*_-_*_i_*: min-max=0.9–3), (2) population median absolute deviation images (*MAD-i*: min-max=0–1), (3) corresponding example participant images (tau: min-max=0.9–2.4; Aβ: min-max=0.9–3), and (4) corresponding example participant segmented deviation regions of interest (ROI) (<L95 =dark blue; L95–M=light blue; M–U95=light red; >U95=dark red). **b**, Spaghetti plots of Jaccard similarity values (percentage) for tau and Aβ across segmented deviation ROIs (two-sided one sample Wilcoxon signed-rank test, **Supplemental Table 2**). **c–e**, Mean percentage overlap across participants for tau¬Aβ (min-max=2–4.5) (**c**), Aβ¬tau (min-max=2–10.5) (**d**), and tau∩Aβ (min-max=2–3.5) (**e**) >U95 ROIs in each parcel of the BFpz, projected on medial and lateral brain surfaces. Parcels are derived from the Human Connectome Project Multi-Modal Parcellation-extended atlas^111^. Darker colours indicate greater overlap of >U95 ROIs with a parcel across participants. **f–h**, Spaghetti plots showing the number of voxels (*left*) and mean SUVr (*right*) across tau¬Aβ (**f**), Aβ¬tau (**g**), and tau∩Aβ (**h**) ROIs for each individual (Friedman test followed by two-sided Dunn’s multiple comparisons test, **Supplemental Table 3–5**). Source data in **Source Data** file.

Tau and Aβ may act both independently and synergistically to alter synaptic function^47-50^. We therefore computed the spatial overlap of tau and Aβ pathology within each individual’s four pathological load intervals (i.e., <L95, L95–M, M–U95, >U95) to generate three unique profiles of pathology: regions with increasing tau but not Aβ (i.e., tau¬Aβ), regions with increasing Aβ but not tau (i.e., Aβ¬tau), and regions with increasing co-localized pathology (i.e., tau∩Aβ) (**Fig. 1b**, **Supplemental Table 2**). We then investigated the anatomical distribution and relative concentration of these distinct pathological profiles across individuals, with a focus on regions with high pathology (>U95) that intersect the BF projection zone (BFpz), including the BF, hippocampus, amygdala, and neocortex. Aligning with known early-stage anatomical patterns of pathology^51,52^, individualized profiles of high tau¬Aβ were found most abundantly in medial and lateral temporal regions (**Fig. 1c**), while high Aβ¬tau profiles were enriched in default mode network hubs (**Fig. 1d**). Profiles of high tau∩Aβ most frequently appeared in the posterior cingulate and occipitotemporal cortex (**Fig. 1e**). Consistent with prior normative mapping frameworks^45^, we observed that regions of high and low pathology were spatially smaller than intermediate pathology regions (**Fig. 1f–h**). We also verified that tau-PET uptake selectively increased across tau¬Aβ regions, Aβ-PET uptake selectively increased across Aβ¬tau regions, and that both radiotracers showed progressive increases moving from low to high tau∩Aβ regions (**Fig. 1f–h**, **Supplemental Table 3–5**).

### Cholinergic synaptic adaptation to tau confers cognitive resilience

In healthy older adults at elevated risk for AD, distinct trajectories of cognitive aging may reflect varying degrees of cholinergic synaptic adaptation to tau and/or Aβ pathology. To interrogate this possibility, we leveraged longitudinal neuropsychological data (Repeatable Battery for the Assessment of Neuropsychological Status (RBANS), *Methods*) from the main cohort PREVENT-AD participants spanning up to 10 years (**Table 1**, main cohort (*n*=64); baseline to final visit: median days=2928, range=1815–3655) and generated annualized percent change (APC) scores for each participant (*Methods*). We then adjusted the APC scores for each participant’s age at follow-up, their baseline cognitive performance, and years of education to isolate individual deviations from expected cognitive aging trajectories.

To derive a data-driven threshold for distinguishing unique cognitive trajectory subgroups, we fit a two-component Gaussian mixture model (GMM) to a distribution of adjusted APC scores from an out-of-sample group of PREVENT-AD participants (**Table 1**, validation cohort (*n*=285); baseline to final visit: median days=2609, range=730–3880). The GMM identified an adjusted APC cut-point at -1.09 (**Fig. 2a**). Using this cut-off, we classified the main and validation cohort participants as resilient (adjusted APC ≥ -1.09) or vulnerable (adjusted APC < -1.09). In both cohorts, no significant subgroup differences were observed in baseline neuropsychological testing performance (**Fig. 2b,c**, **Supplemental Table 6,7**) years of formal education, proportion of females versus males, or number of apolipoprotein E ε4 (*APOE* ε4) carriers (**Table 2**).

**Fig. 2.**
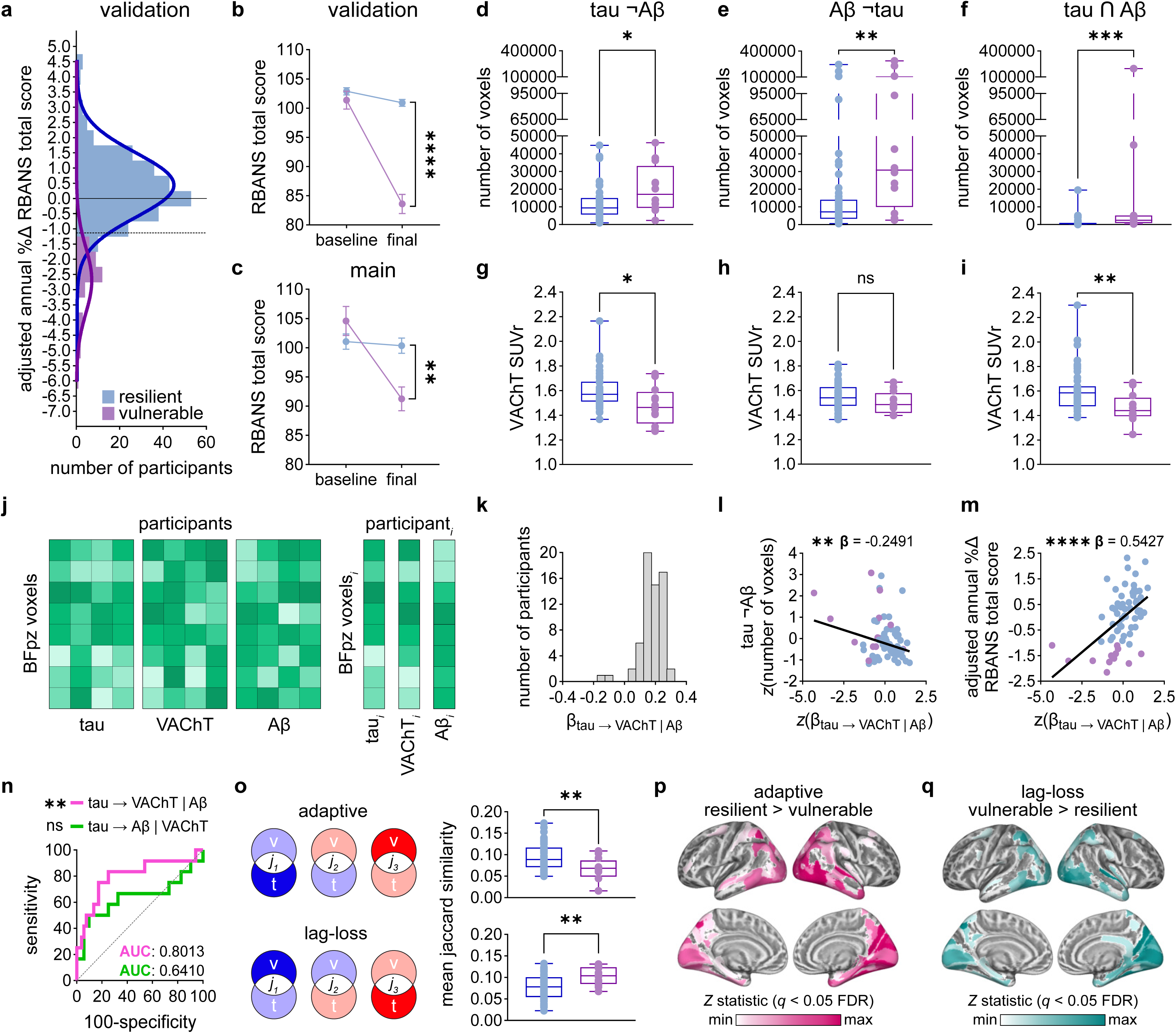
Cholinergic synaptic adaptation to tau confers cognitive resilience. **a**, Resilient (*n*=245) and vulnerable (*n*=40) subgroups in the validation cohort of PREVENT-AD participants, identified from multivariate Gaussian mixture modelling (GMM) of adjusted RBANS annualized percent change (APC) scores. The dashed line indicates the GMM-derived cut-point for subgroup classification and the solid line denotes an APC=0 (no change). **b,c**, RBANS total scores at baseline and final visits for validation (**b**) and main (**c**) PREVENT-AD cohorts, split by subgroup (mean±s.e.m.; repeated mixed-effects model followed by two-sided Fisher’s LSD test, **Supplemental Table 6,7**). **d–f**, Box-and-whisker plots showing the number of voxels in high (>U95) tau¬Aβ (**d**), Aβ¬tau (**e**), tau∩Aβ (**f**) regions for resilient (blue) and vulnerable (purple) subgroups in the main PREVENT-AD cohort (two-sided Mann-Whitney *U* test). **g–i**, Box-and-whisker plots showing VAChT standardized uptake value ratio (SUVr) levels in the above regions (two-sided Mann-Whitney *U* test). **j**, Schematic of within-participant regression analyses. For each participant, individual (*i*) voxel-wise profiles of tau-PET, VAChT-PET, and Aβ-PET within the BFpz were used to compute the linear relationship between tau and VAChT, while covarying for Aβ. **k**, Histogram of individual tau-VAChT regression coefficients (β). **l**, Scatter plot showing the robust linear association between tau-VAChT coupling coefficients from **k** and the number of voxels in high (>U95) tau¬Aβ regions. **m**, Scatter plot showing the robust linear association between tau-VAChT coupling coefficients from **k** and adjusted APC RBANS total score values. **n**, Receiver-operating curves (ROCs) generated using within-participant regression coefficients to classify resilient and vulnerable subgroups. **o**, Schematic of adaptive (*top*) and lag-loss (*bottom*) models of spatial overlap between tau and VAChT deviation intervals. Box-and-whisker plots showing mean Jaccard similarity values within participants from each model, split by resilient and vulnerable subgroup (two-sided Mann-Whitney *U* test). **p,q**, Subgroup comparisons of the mean overlap of voxel-level Jaccard similarity values with parcels in the BFpz for adaptive (**p**, *Z* min-max=2.39–4.55) and lag-loss (**q**, *Z* min-max= 2.52–3.84) models, displayed on medial and lateral brain surfaces. Parcels are derived from the Human Connectome Project Multi-Modal Parcellation-extended atlas^111^ (two-sided Wilcoxon rank-sum test; adjusted for the false-discovery rate (FDR) at 5%, **Supplemental Table 8,9**). For box-and-whisker plots, the center line indicates the median, hinges reflect the interquartile range (IQR), whiskers extend to the minimum and maximum values, and each dot represents an individual participant. \**P*<0.05, \*\**P*<0.01, \*\*\**P*<0.001, \*\*\*\**P*<0.0001, ns, not significant. Source data in **Source Data** file.

**Table 2.**
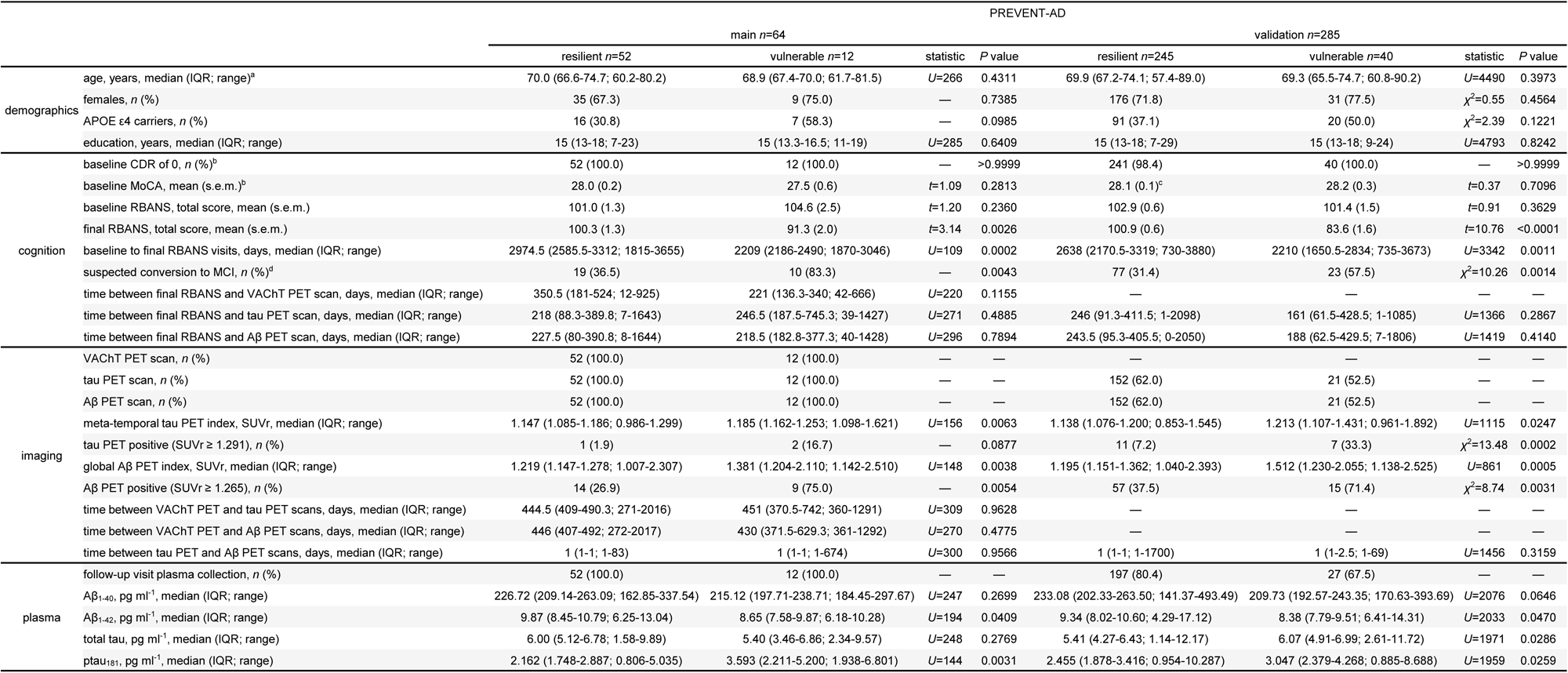
PREVENT-AD resilient and vulnerable subgroup demographic comparisons. ^a^age at final RBANS visit. ^b^information on inclusion criteria for individuals with a baseline CDR > 0 or MoCA < 26 can be found in *Methods*. ^c^missing MoCA data for one resilient participant, mean (standard error of the mean) and corresponding statistics performed for *n*=244. ^d^suspected conversion status as of 2025; information on evaluation of clinical progression can be found in *Methods*. Continuous data are presented as the mean (standard error of the mean) or median (interquartile range; range). Categorical data are presented as the number of participants (percentage). Differences in continuous data were assessed using the two-sided independent samples *t* test or two-sided Mann–Whitney *U* test, as appropriate. The two-sided Fisher’s exact test and the Chi-squared test were used to assess differences in categorical variables, as appropriate. Exact *P* values are reported. Aβ, β-amyloid; APOE ε4, apolipoprotein E ε4 allele; CDR, clinical dementia rating; MCI, mild cognitive impairment; MoCA, Montreal cognitive assessment; ptau_181_, phosphorylated tau at threonine-181; RBANS, repeatable battery for the assessment of neuropsychological status; SUVr, standardized uptake value ratio; VAChT, vesicular acetylcholine transporter.

We next asked whether the resilient and vulnerable subgroups in the main cohort differed according to their individual profiles of tau and/or Aβ. To do this, we compared the size of the high (>U95) tau¬Aβ, Aβ¬tau, and tau∩Aβ regions between groups^53,54^. Consistent with an increased spatial extent of pathology, the vulnerable subgroup exhibited significantly larger tau¬Aβ (**Fig. 2d**, *U*=189, *P*=0.0339), Aβ¬tau (**Fig. 2e**, *U*=149, *P*=0.0042), and tau∩Aβ (**Fig. 2f**, *U*=99, *P*=0.0001) regions than the resilient group. We then compared VAChT levels between resilient and vulnerable groups in each of these pathophysiologically distinct regions. Here, we observed that the resilient subgroup exhibited significantly higher VAChT levels in tau¬Aβ (**Fig. 2g**, *U*=168, *P*=0.0122) and tau∩Aβ (**Fig. 2i**, *U*=153, *P*=0.0064) regions relative to vulnerable individuals. However, VAChT levels did not significantly differ between subgroups in areas defined only by high Aβ pathology (**Fig. 2h**, *U*=232, *P*=0.1737). These findings suggest that tau may more directly affect cholinergic presynaptic function than Aβ.

Do the observed group differences in VAChT reflect tau-associated cholinergic presynaptic loss in vulnerability, tau-mediated VAChT upregulation in resilience, or both? To investigate these possibilities, we performed voxel-wise regressions between each individual’s profile of tau and VAChT in the BFpz, while covarying for Aβ (**Fig. 2j**, *Methods*). This approach allowed us to isolate fine-grained co-fluctuations in the spatial relationship between tau and VAChT. We found that tau-mediated changes in VAChT varied in both direction and magnitude across the cohort (**Fig. 2k**), with a skew toward positive associations (median *β*=0.1838, range=-0.1561–0.2909), consistent with an upregulation of VAChT to tau.

To confirm that the observed increases in [^18^F]-FEOBV uptake in humans reflect endogenous increases in presynaptic VAChT levels, we performed [^18^F]-FEOBV microPET imaging in a *BAC*-transgenic mouse line designed to globally overexpress presynaptic VAChT (*hyp*)^55^ and wild-type (*wt*) controls (**Extended Data Fig. 5a**). We found that *hyp* mice exhibited on average ∼20% higher [^18^F]-FEOBV uptake throughout the BFpz relative to *wt* mice (*hyp* median V_T_=1.619; *wt* median V_T_=1.373, **Extended Data Fig. 5b**). In a subset of these mice, we further demonstrated that [^18^F]-FEOBV uptake closely tracked with VAChT protein levels (**Extended Data Fig. 5c**) in the cortex (**Extended Data Fig. 5d**, *ρ*=0.8857, *P*=0.0167) and hippocampus (**Extended Data Fig. 5d**, *ρ*=0.7714, *P*=0.0514). In humans, [^18^F]-FEOBV uptake increased by ∼4.7% per standard deviation increase in tau, a response magnitude well within the dynamic range observed in VAChT overexpressing mice.

If tau-colocalized upregulation of presynaptic VAChT reflects a functionally adaptive response, it should be associated with smaller spatial extents of tau pathology in the BFpz and more favourable cognitive trajectories. Consistent with this hypothesis, robust linear regression analysis revealed that stronger positive tau-VAChT coupling was linked to reduced spatial coverage of high (>U95) tau¬Aβ region (**Fig. 2l**, *β*[*SE*]=-0.2491[0.0929], *P*=0.0094) and more stable longitudinal cognition scores (**Fig. 2m**, *β*[*SE*]=0.5427[0.1215], *P*=0.00003) in healthy older adults at elevated risk for AD. We further examined whether the time between tau-PET and VAChT-PET data acquisition influenced the observed associations in main cohort participants (**Table 1**). After adjusting for the tau-to-VAChT PET interscan interval, the tau-VAChT coupling coefficients remained significantly negatively associated with the spatial extent of high tau¬Aβ regions (*β*[*SE*]=-0.2593[0.0990], *P*=0.0111) and significantly positively associated with adjusted APC values (*β*[*SE*]=0.5347[0.1229], *P*=0.00005). We also observed no significant association between tau-to-VAChT PET interscan intervals and tau-VAChT coupling coefficients (*β*[*SE*]=-0.1146[0.1010], *P*=0.2606).

We then evaluated the ability of tau-VAChT coupling to discriminate resilient and vulnerable subgroups. Here, we found that individual tau-VAChT coupling coefficients effectively distinguished the groups with an area under the curve (AUC) of 0.8013 (**Fig. 2n**, 95CI=0.6411–0.9615, *P*=0.0012). At this threshold, tau-VAChT coupling corresponds to a ∼1.1% increase in VAChT per standard deviation increase in tau. As a comparison, we tested whether direct coupling between tau and Aβ while adjusting for VAChT could also determine subgroup membership. However, this model produced a substantially lower, nonsignificant AUC of 0.6410 (**Fig. 2n**, 95CI=0.4235–0.8585, *P*=0.1301). These results suggest that the relationship between tau and VAChT, rather than tau and Aβ, more strongly distinguishes individual cognitive aging trajectories in healthy older adults at risk for AD.

Although the linear relationship between VAChT and tau in the BFpz discriminates resilient and vulnerable subgroups (**Fig. 2n**), it does not capture regions where VAChT is disproportionately higher or lower relative to levels of colocalized tau. We therefore examined whether VAChT and tau are also spatially colocalized when their deviation profiles are offset relative to one another. To enable direct comparison of VAChT and tau deviation intervals within participants, we applied the normative deviation mapping approach to the VAChT-PET dataset, yielding stratified cholinergic presynaptic profiles for each individual (*Methods*, **Extended Data Fig. 4**, **Supplemental Table 1**). We then compared the spatial overlap between adjacent VAChT and tau deviation intervals under two biologically motivated models of alignment: an *adaptive* model, in which VAChT occupies a deviation interval one level higher than tau, and a *lag-loss* model, in which tau occupies a deviation interval one level higher than VAChT (**Fig. 2o**). Here, we found that resilient individuals exhibited significantly greater overlap under the adaptive model (**Fig. 2o**, *U*=162, *P*=0.0089), whereas vulnerable individuals showed significantly greater overlap under the lag-loss model (**Fig. 2o**, *U*=161, *P*=0.0083). We then examined where in the brain these distinct models are most strongly expressed. At the group level, both models exhibited highest overlap in temporal and precuneus regions, indicating a common core brain network expressing strong tau-VAChT colocalization (**Extended Data Fig. 6a,b**). However, resilient and vulnerable individuals showed opposite load level-dependent profiles of VAChT and tau within these areas (**Fig. 2p,q**). Together, these findings suggest that where presynaptic VAChT and tau colocalize, VAChT levels tend to overshoot tau in the resilient brain, and undershoot tau in the vulnerable brain.

### Cholinergic neurons are enriched for synaptic plasticity

The spatially coordinated, tau-sensitive cholinergic presynaptic response suggests an underlying transcriptional program tuned to both presynaptic plasticity and tau pathology. To explore the cellular basis of this mechanism, we analyzed single-nucleus RNA sequencing (snRNAseq) atlases from the healthy adult human brain^56^ and healthy adult mouse brain^57^. Low-dimensional embedding of these datasets revealed a conserved architecture across the telencephalic landscape, comprising 235 and 66 neuronal clusters in the human and mouse datasets, respectively. Neuronal clusters were further annotated according to their primary anatomical origin across the telencephalic BFpz, including the BF, amygdala, cortex, and hippocampus (**Fig. 3a**), as well as their dominant neurochemical identity (i.e., glutamatergic, GABAergic, cholinergic) (**Fig. 3b**).

**Fig. 3.**
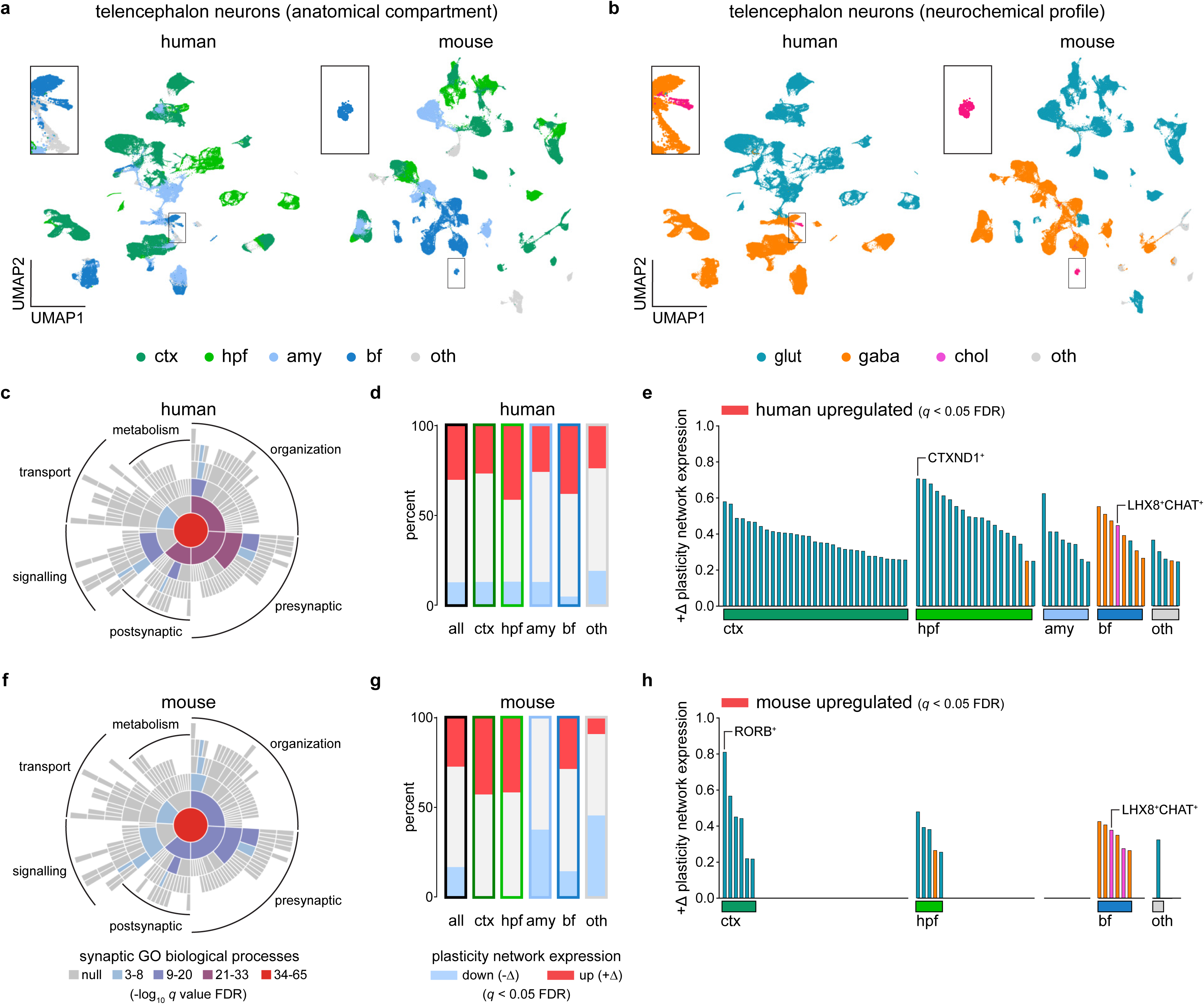
Plasticity and vulnerability converge in basal forebrain cholinergic neurons and their targets. **a,b**, UMAP embedding of telencephalon neurons from the health adult human^56^ and adult mouse^57^ brain, colour-coded by primary telencephalic compartment (**a**) and major neurochemical class (**b**). **c,f**, Sunburst plots of enriched *SynGO*^59^ biological processes according to plasticity network genes^58^ overlapping with the human (**c**) or mouse (**f**) snRNAseq datasets, colour-coded by enrichment *q* value (one-sided Fisher’s exact test; adjusted for the false-discovery rate (FDR) at 1%). **d,g**, Stacked bar plots showing the percentage of human (**d**) or mouse (**g**) telencephalic neuronal clusters exhibiting significant upregulation (+Δ) or downregulation (-Δ) of the plasticity gene-network (permutation testing; adjusted for the FDR at 5% across clusters, **Supplemental Table 10,11**). **e,h**, Bar plots displaying neuronal clusters in the human (**e**) or mouse (**h**) telencephalon with significant positive plasticity gene-network module scores, grouped according to their primary anatomical compartment and colour-coded by dominant neurochemical profile. Source data in **Source Data** file.

We then systematically examined whether an *a priori* gene-network linked to both tau pathology and presynaptic plasticity in vulnerable cortical regions^58^ is also highly expressed in BF cholinergic neurons. We first confirmed that the overlap of genes between the plasticity gene-network and the human (**Fig. 3c**) and mouse (**Fig. 3f**) snRNAseq datasets were functionally enriched for presynaptic biological processes, using an independently curated synaptic gene set^59^. Next, we quantified plasticity gene-network expression across all telencephalic neuronal clusters in the human and mouse snRNAseq atlases (*Methods*). Mapping plasticity gene-network enrichment to each of the neuronal clusters revealed a unique distribution of this transcriptional program across anatomical compartments in both species (**Fig. 3d,g**, **Supplemental Table 10,11**), with 38% and 28% of neuronal clusters within the human and mouse BF, respectively, showing significant upregulation. Consistent with this pattern, human (**Fig. 3e**, Δ=0.4488) and mouse (**Fig. 3h**, Δ=0.3799) BF cholinergic neuronal clusters (i.e., *LHX8^+^CHAT^+^*) exhibited significant positive enrichment for the plasticity gene-network (FDR < 0.05). Other telencephalic neurons positively enriched for the plasticity gene-network included *CTXND1^+^* glutamatergic hippocampal neurons in humans (**Fig. 3e**, Δ=0.7081) and *RORB*^+^ glutamatergic upper-layer cortical neurons in mice (**Fig. 3h**, Δ= 0.8109). Ranking across all telencephalic neuronal clusters further revealed that BF cholinergic neurons were among the highest expressors of the plasticity gene-network in both species (**Fig. 3e**, 24^th^ of 235; **Fig. 3h**, 10^th^ of 66), consistent with the hypothesis that BF cholinergic neurons are transcriptionally primed to support presynaptic plasticity programs linked to tau vulnerability.

To determine whether specific components of the plasticity gene-network were selectively upregulated in BF cholinergic neurons, we also performed gene-level enrichment analyses comparing BF cholinergic neurons to the other neuronal clusters of the telencephalon (*Methods*). Functional annotation of the genes significantly upregulated in BF cholinergic neurons revealed strong enrichment for microtubule-based processes in the human dataset, and localization to the microtubule cytoskeleton in both species (FDR < 0.05). Notably, these enriched gene sets included the microtubule-associated protein tau (*MAPT*) gene. Together, these findings suggest that microtubule cytoskeletal regulation represents a central component of the plasticity gene-network expressed in BF cholinergic neurons.

### Cholinergic synaptic function is required for cortical plasticity

The human imaging and snRNAseq findings suggest that BF cholinergic neurons are endowed with intrinsically high expression of a gene-network supporting presynaptic plasticity. We therefore causally probed whether disruption of cholinergic presynaptic function impairs cortical plasticity. To do this, we leveraged a conditional forebrain VAChT knockout (*ko*) mouse model^28,60,61^ and examined how loss of VAChT-mediated functions impacts reversal learning and structural integrity.

In *ko* mice, VAChT protein expression is selectively eliminated from forebrain cholinergic neurons through *Cre*-dependent recombination^61^. Biochemical analyses confirmed that VAChT protein levels were severely reduced in forebrain VAChT *ko* mice relative to *wt* mice in cortical and hippocampal targets of the cholinergic BF, as well as striatal cholinergic interneurons (**Fig. 4a–c**, *U*=0, *P*=0.0286), whereas regions innervated by brainstem cholinergic neurons remained intact (**Fig. 4d**, *U*=6, *P*=0.6857). To extend these findings *in vivo*, we performed [^18^F]-FEOBV microPET imaging in both forebrain VAChT *ko* and *wt* mice (**Fig. 4e**). Consistent with the selective elimination of VAChT in forebrain cholinergic neurons, forebrain VAChT *ko* mice exhibited reduced forebrain [^18^F]-FEOBV uptake relative to *wt* mice (**Fig. 4f**, *U*=16, *P*=0.0441), with no corresponding differences observed in the brainstem (**Fig. 4f**, *U*=38, *P*>0.9999).

**Fig. 4.**
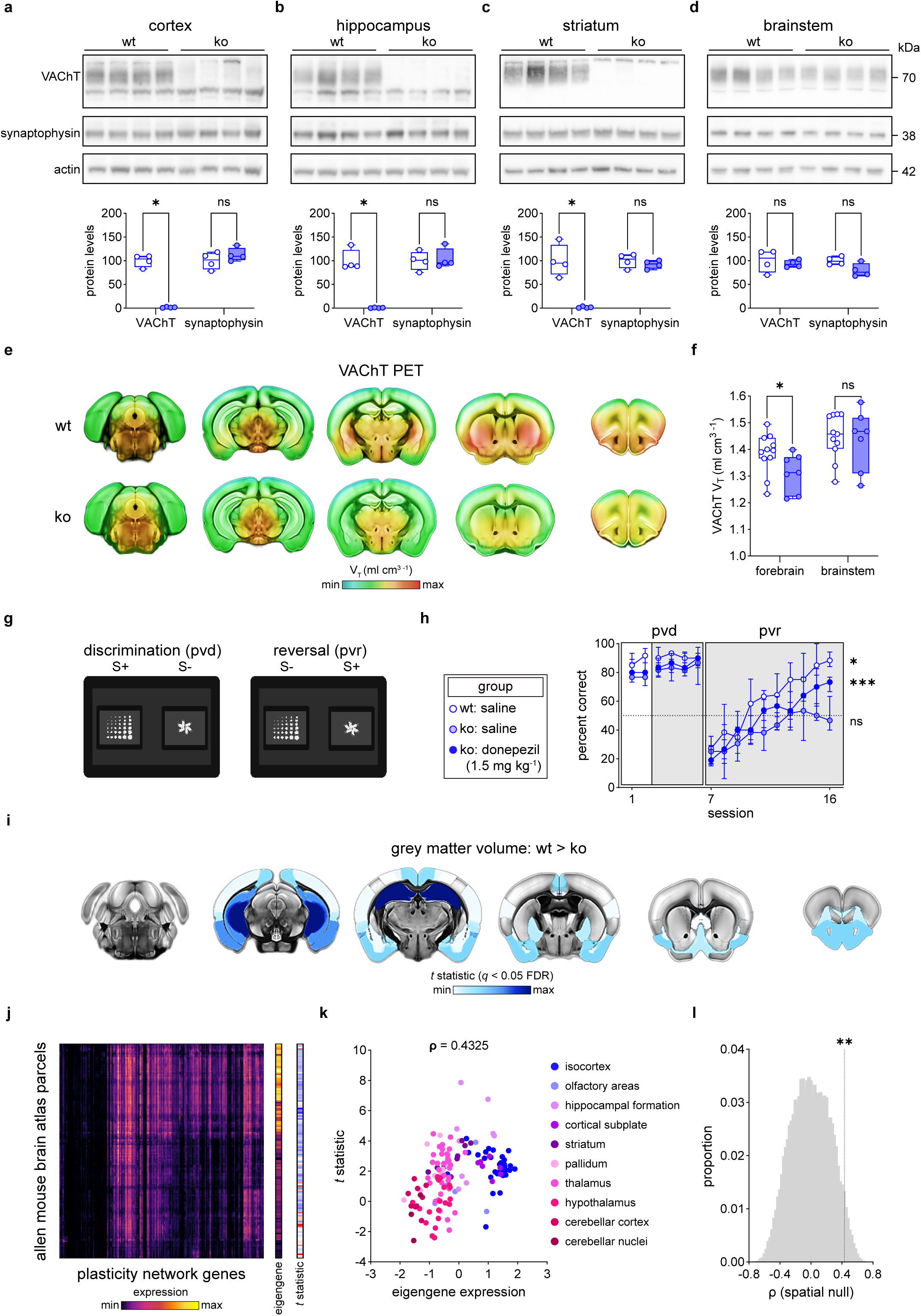
Loss of forebrain VAChT impairs cognitive flexibility and induces hippocampal neurodegeneration in mice. **a–d**, Immunoblot analysis of VAChT and synaptophysin protein levels in *wt* (*n*=4) and forebrain VAChT *ko* (*n*=4) mice from cortical (**a**), hippocampal (**b**), striatal (**c**), and brainstem (**d**) tissue samples. In all regions, synaptophysin protein levels did not differ between genotypes (**Supplemental Table 12–15**). Synaptophysin and actin were used as protein loading controls (two-sided Mann-Whitney *U* test). **e**, Average [^18^F]-FEOBV total volume of distribution (V_T_) brain profiles in *wt* (*n*=11) and forebrain VAChT *ko* (*n*=7) mice, overlaid on coronal sections of the mouse brain (min-max= 0.3–2.1). **f**, Forebrain and brainstem [^18^F]-FEOBV V_T_ values in *wt* and forebrain VAChT *ko* mice (two-sided Mann-Whitney *U* test). **g**, Schematic of the mouse pairwise visual discrimination-reversal (*pvd*–*pvr*) touchscreen task. **h**, Plot showing *pvd*–*pvr* task performance (percentage of correct trials) in saline-treated *wt* mice (*n*=6), saline-treated forebrain VAChT *ko* mice (*n*=8), and donepezil-treated forebrain VAChT *ko* mice (*n*=11). Sessions 1–6 are *pvd*, and sessions 7–16 are *pvr*. The grey background indicates the treatment-on sessions, and the horizontal dotted line indicates chance (50%) performance (median±IQR; one-sided one sample Wilcoxon signed-rank test, **Supplemental Table 16**). **i**, Between-group comparisons of regional grey matter volume in the BFpz (*wt*: *n*=14, forebrain VACHT *ko*: *n*=10). Regions from the Allen Mouse Brain Atlas common coordinate framework version 3^125^ overlaid on coronal sections of the mouse brain, thresholded and colour-coded by *t* statistic (*t* min-max=2.35–6.85; two-sided independent samples *t* test; adjusted for the false-discovery rate (FDR) at 5%, **Supplemental Table 17**). **j**, (*left*) Heatmap displaying expression levels of plasticity network genes^58^ across parcels from the Allen Mouse Brain Atlas^65^, ordered using hierarchical clustering (expression min-max: 0–50). (*right*) Eigengene and *t* statistic values for each parcel in the Allen Mouse Brain Atlas (**Supplemental Table 19**). **k**, Scatterplot showing the relationship between the spatial profile of grey matter volume differences in *wt* versus forebrain VAChT *ko* mice, and the eigengene of plasticity gene-network expression, colour-coded according to each parcel’s region of origin (two-sided Spearman rank correlation). **l**, Histogram showing the distribution of 40,000 surrogate null Spearman ρ correlations computed from spatially autocorrelated random maps. The dashed line marks the observed correlation, and the asterisks indicate the corresponding empirical *P* value. For box-and-whisker plots, the center line indicates the median, hinges reflect the interquartile range (IQR), whiskers extend to the minimum and maximum values, and each dot represents an individual animal. Full membranes for **a–d** are displayed in **Supplemental Fig. 2**. \**P*<0.05, \*\**P*<0.01, \*\*\**P*<0.001, ns, not significant. Source data in **Source Data** file.

To assess reversal learning under normal or reduced forebrain VAChT conditions, *wt* and forebrain VAChT *ko* mice performed a pairwise visual discrimination and reversal (*pvd*–*pvr*) touchscreen task of cognitive-flexibility (**Fig. 4g**, *Methods*). During *pvd* testing, both *wt* and forebrain VAChT *ko* mice performed significantly above change (>50 percent correct) at target detection (**Fig. 4h**, **Supplemental Table 16**). However, consistent with a causal role for forebrain cholinergic synaptic plasticity in updating learned associations^62-64^, forebrain VAChT *ko* mice exhibited a sustained impairment during reversal sessions (**Fig. 4h**, session 16: *W*=15, *P*=0.4609). By contrast, *wt* mice successfully achieved reversal learning (**Fig. 4h**, session 16: *W*=21, *P*=0.0156). We further demonstrated that partial restoration of acetylcholine bioavailability in forebrain VAChT *ko* mice via continuous administration of an acetylcholinesterase inhibitor (donepezil; 1.5 mg kg^-1^) significantly improved *pvr* task performance to above chance levels (**Fig. 4h**, session 16: *W*=66, *P*=0.0005).

In humans, *in vivo* measures of BF grey matter volume loss track the progression of cortical degeneration across the AD continuum^19^. Notably, the spatial pattern of cortical atrophy mirrors the topography of presynaptic cholinergic deficits observed in late disease stages^21^. We therefore hypothesized that a targeted distruption of forebrain cholinergic presynaptic function may compromise the structural integrity of the BFpz. To investigate this possibility, we performed whole-brain *in vivo* 9.4T structural magnetic resonance imaging (MRI) in forebrain VAChT *ko* and *wt* mice and applied voxel-based morphometry to quantify grey matter volume differences between the groups. Compared to *wt* mice, we observed significantly lower grey matter volume across multiple BFpz regions in forebrain VAChT *ko* mice, with the most pronounced reductions observed in the hippocampus (**Fig. 4i**, **Supplemental Table 17**). We additionally confirmed in age-matched *Cre-*only expressing mice that the observed changes in hippocampal structural integrity were not due to off-target recombination events (**Supplemental Table 18**). Together, these findings further reinforce the critical contributions of BF cholinergic presynaptic function to the mechanisms supporting cortical plasticity.

We noted that the spatial pattern of grey matter volume loss in forebrain VAChT *ko* mice overlapped with regions identified in the snRNAseq analysis as enriched for plasticity. To test this overlap formally, we derived an eigengene representation of the plasticity gene-network across parcels of the Allen Mouse Brain Atlas^65,66^ and examined its spatial correlation with grey matter volume differences in the same atlas space (**Fig. 4j**, **Supplemental Table 19**). Here, we observed that regions showing greater grey matter volume loss in forebrain VAChT *ko* mice exhibited higher intrinsic expression of the plasticity gene-network (**Fig. 4k**, *ρ*=0.4325), an association that significantly exceeded chance under spatially constrained null models^67^ (**Fig. 4l**, empirical *P*=0.0097). Hence, neurons transcriptionally primed for high plasticity appear to converge on those vulnerable to degeneration under conditions of reduced forebrain VAChT expression.

## Discussion

We demonstrate that synaptic plasticity serves as a common mechanism through which BF cholinergic neurons both adapt and contribute to early tau dysregulation, increasing their vulnerability to disease. Under physiological conditions, VAChT regulates cholinergic synaptic output in response to local network activity, while a *MAPT*-associated gene-network supports the presynaptic structural and molecular adaptations required to sustain this plasticity. Among the at-risk older adults studied here, our identification of both cognitively resilient and vulnerable subgroups suggests that there are distinct profiles of cholinergic synaptic adaptation to tau pathology in the aging brain. In resilient individuals, tau-induced synaptotoxicity was accompanied by a robust presynaptic VAChT response. In vulnerable individuals, a reduced capacity to mount this adaptive response was associated with a greater spatial extent of tau pathology in the brain.

The fundamental relationship between cholinergic presynaptic function and tau aligns with theoretical models of homeostatic synaptic plasticity^8-11,68^, which posit that neurons adaptively regulate synaptic efficacy to preserve stability under sustained perturbation. Our data offers an *in vivo* demonstration of this mechanism in humans, wherein a graded presynaptic cholinergic response to tau pathology discriminates distinct cognitive trajectories and is associated with the spatial extent of tau pathology in the aging brain. Importantly, the failure of homeostatic synaptic plasticity in some individuals may explain the inflection point between normal brain aging and the AD trajectory. Such a progression echoes broader models of cellular aging, in which early transcriptomic, epigenetic, or metabolic disruptions trigger compensatory cellular adaptations that ultimately fail, leading to disease phenotypes^69^.

Although we cannot exclude the contribution of soluble or intracellular Aβ to early changes in cholinergic presynaptic function^70-75^, particularly during the initial stages of AD, the insoluble Aβ pathology quantified by Aβ-PET^76^ in this study did not colocalize with adaptive VAChT upregulation. Recent studies combining tau-PET and Aβ-PET imaging with cortical gene expression profiles^77,78^ have shown that these proteinopathies propagate along distinct transcriptional pathways, with tau propagation aligning to axon-related genes, including *MAPT*, and Aβ propagation associating with dendrite-related profiles. These findings, together with our observations, suggest that while Aβ-related mechanisms may contribute to early synaptic dysfunction, the transcriptional programs governing tau-mediated cholinergic presynaptic adaptation and vulnerability operate through at least partially independent pathways.

Our findings in forebrain VAChT *ko* mice demonstrate causal interrelationships among cholinergic signalling, synaptic plasticity and the structural integrity of the BFpz. When the dynamic range of cholinergic neurotransmission is chronically constrained, as in forebrain VAChT *ko* mice^79^, plasticity-dependent reversal learning is impaired. These deficits are partially rescued by enhancing acetylcholine availability, further underscoring the functional dependence of cognitive flexibility on forebrain cholinergic presynaptic plasticity^62-64^. Forebrain VAChT *ko* mice also exhibited reductions in grey matter volume in brain regions densely innervated by BF cholinergic projections, including the hippocampus. We found that this profile of neurodegeneration spatially covaried with regional enrichment for a gene-network supporting plasticity^58^ in the healthy adult mouse brain. Deprivation of VAChT in cholinergic presynaptic terminals may therefore disrupt cholinergic surveillance of transcriptional programs involved in plasticity, including tau proteostasis, initiating a cascade of cellular dysfunctions resembling early AD pathology^28,60,61^.

Although *MAPT* is traditionally associated with microtubule stabilization, it exhibits strong co-expression with genes involved in presynaptic plasticity, vesicular transport, and neurotransmitter release^58,80,81^. Complementary experimental evidence suggests that one of the primary loci of tau pathology is in presynaptic terminals^82,83^, where its mislocalization disrupts axonal transport, neurotransmitter release probability, and vesicle recycling through aberrant interactions with synaptic proteins and trafficking complexes. These dysfunctions are exacerbated by the unique morphofunctional properties of vulnerable neurons, including highly branched, lightly myelinated axons^84^, and a sustained propensity for activity-dependent remodelling in adulthood^85^. Using snRNAseq atlases to survey neurons in the healthy adult human^56^ and mouse^57^ telencephalon, we show that an *a priori* plasticity gene-network^58^ is highly enriched in multiple subtypes of neurons which populate vulnerable brain regions, including BF cholinergic and GABAergic neurons and glutamatergic projection neurons of the hippocampus, cortex and amygdala. Together, our findings support the hypothesis that intrinsic, plasticity-linked transcriptional profiles predispose specific neuron types to early tau dysregulation^85-89^. Consistent with this idea, prior neuropathological work has shown that tau neurofibrillary tangles are present in virtually all older adults’ brains by the age of 60^18,90^, and tend to colocalize areas densely innervated by BF cholinergic projections^85,87,91^.

Several limitations of the present study warrant consideration. The overall small sample size (*n*=64) of participants with VAChT-PET, tau-PET, and Aβ-PET data underscores the need for replication in larger, independent studies. However, the resilient and vulnerable classification approach employed here was derived from a substantially larger sample (*n*=285) of longitudinal cognitive data in the broader PREVENT-AD cohort, supporting the stability of this subgrouping strategy. Another limitation is the cross-sectional nature of the triple PET radiotracer dataset, which precludes inferences regarding the temporal sequence of cholinergic presynaptic adaptations relative to Aβ and tau progression. Longitudinal imaging in presymptomatic, at-risk older adults will be necessary to clarify these interrelationships. Finally, the participants in the present study were approximately 70 years old during the PET imaging period. The results of the present work therefore likely do not capture prior interactions between cholinergic presynaptic function and pathophysiological processes that may arise in midlife. Extending similar molecular imaging paradigms to younger populations will be necessary for mapping early alterations in cholinergic presynaptic plasticity and evaluating its utility for discriminating resilient and vulnerable cognitive aging trajectories.

The link between VAChT and tau helps articulate new directions for future research. Key priorities include integrating *in vivo* biomarkers for synaptic function and density alongside biomarkers of AD pathology in longitudinal studies of aging and at-risk populations^50,92-95^. In parallel, greater consideration should be given to the development of novel treatment approaches aimed at promoting plasticity-dependent cognitive processes to strengthen brain resilience^96,97^. Our findings also raise intriguing possibilities. Does the adaptive cholinergic presynaptic response exist in other neurodegenerative disorders^98-100^? Why does this response fail in some before others? Can this mechanism be pharmacologically modulated^101^? Answers to these questions may inform therapeutic strategies that complement monoclonal antibodies targeting Aβ or tau^102^.

## Methods

### Human Ethics Statement

All participants provided written informed consent at baseline and all follow-up visits. The consent forms along with study protocols for the data acquisition procedures (i.e., imaging, neuropsychological testing) were approved by the Institutional Review Board at McGill University and the CIUSSS de l’Ouest de l’Ile de Montréal Research Ethics Board (Montréal, QC, Canada).

### Participant Eligibility and Enrollment

Participants were recruited from the PRe-symptomatic EValuation of Experimental or Novel Treatments for Alzheimer’s Disease (PREVENT-AD) program (preventad.loris.ca). Inclusion and exclusion criteria for the PREVENT-AD program were as reported previously^33,34^. In brief, eligibility for enrollment in the PREVENT-AD research program requires a parental history of sporadic Alzheimer-like disease or at least two affected siblings. Participants were also required to be ≥60 years of age, or 55–59 years if within 15 years of the age at symptom onset of their affected first-degree relative. At the screening visit, the Montreal Cognitive Assessment (MoCA)^103^ and the Clinical Dementia Rating (CDR)^104^ test were administered to exclude cognitive impairment. Individuals with a MoCA score < 26 or a CDR > 0 were only enrolled if a comprehensive neuropsychological evaluation performed by a clinician confirmed intact cognition. The Repeatable Battery for the Assessment of Neuropsychological Status (RBANS) was also administered at baseline, and participants scoring below age-adjusted normative values underwent further comprehensive evaluation to confirm normal cognitive status.

At baseline, participants’ biological sex and total years of formal education were recorded via self-report. Apolipoprotein E (*APOE*) genotyping was also performed using DNA isolated from 200 µl of blood collected. DNA extraction was carried out with the QIASymphony system and QIAamp DNA Blood Mini Kit (Qiagen, Valencia, CA, USA), and genotyping was conducted by pyrosequencing on the PyroMark Q96 platform (Qiagen, Toronto, ON, Canada).

### Clinical Progression

To evaluate clinical progression in PREVENT-AD participants (**Table 1,2**), cognitive data were reviewed in a multi-step process blind to genetic, imaging, and biomarker data^34^. Suspected mild cognitive impairment (MCI) cases were initially identified by trained research personnel, after which all available longitudinal cognitive assessments were independently examined by two neuropsychologists. Participants meeting criteria for suspected MCI were subsequently adjudicated at a multidisciplinary consensus meeting held in 2025 by a cognitive neurologist, a neuropsychiatrist, and the two reviewing neuropsychologists. Final classification as cognitively unimpaired or suspected MCI was determined based on the entirety of available longitudinal data and remained blinded to biomarker information. As previously reported^34^, the majority of PREVENT-AD participants classified as suspected MCI have a CDR score of 0. Thus, PREVENT-AD participants identified as suspected MCI present with relatively subtle cognitive deficits, potentially corresponding to cases that might be characterized as subjective cognitive decline in other research cohorts or in a memory clinic setting.

### RBANS and Adjusted Annualized Percent Change Scores

Participants completed Repeatable Battery for the Assessment of Neuropsychological Status (RBANS) evaluations at baseline and follow-up visits^105^. The RBANS consists of 12 subtests (i.e., list learning, story memory, figure copy, line orientation, picture naming, semantic fluency, digit span, coding, list recall, list recognition, story recall, and figure recall), which contribute to five index scores (i.e., immediate memory, delayed memory, language, attention, and visuospatial construction), as well as a composite total score. To mitigate practice effects, equivalent alternate forms of the assessment battery were administered across study visits.

For the present study, all participants were required to have a minimum of two years between their baseline and final RBANS evaluations. A total of 349 PREVENT-AD participants met this criterion. Of these, 64 participants were in the main cohort, and thus the remaining 285 were assigned to the validation cohort. **Table 1** provides a summary of the intervals between baseline and final RBANS evaluations for both cohorts.

Longitudinal cognitive change was indexed using an annualized percent change (APC) metric derived from RBANS scores (total or index):

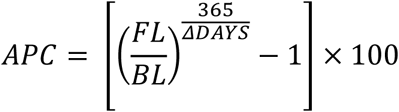

where *BL* and *FL* refer to baseline and final RBANS scores (total or index), respectively, and *ΔDAYS* represents the number of days elapsed between those two visits. The APC calculation accounts for inter-individual variability in follow-up duration and quantifies yearly change in cognitive performance^101^. For main cohort participants, the final RBANS visit was defined as that which was within the triple PET radiotracer imaging period (**Extended Data Fig. 1**). For 11 participants whose VAChT-PET scans were acquired shortly before the COVID-19 pandemic (February 2020), the final RBANS visit was defined as the next available neuropsychological assessment, conducted when testing resumed in 2021-2022. For validation cohort participants, the final RBANS visit was defined as the last recorded assessment in *data_release_8.0* (https://openpreventad.loris.ca).

Using these data, adjusted APC scores were computed using a linear regression model with age at final RBANS visit, baseline RBANS performance, and years of education as predictors. The residuals of this model represent deviations from expected cognitive trajectories, independent of demographic and baseline cognitive performance.

To identify latent subgroups of cognitive aging, Gaussian mixture modelling (GMM) was applied to a multivariate representation of cognitive change. Specifically, adjusted APC scores from all five RBANS index scores and the composite total score were included in the model. The GMM was applied to the adjusted APC scores from the validation cohort (*n*=285) using maximum likelihood estimation, initialized with 50 random starts and 10,000 maximum iterations to ensure robust convergence. This model revealed a bimodal structure in the adjusted APC distribution, and a threshold of -1.09 was derived from the posterior probabilities to optimally distinguish declining (i.e., vulnerable) versus stable (i.e., resilient) cognitive trajectories. The cut-off derived from the validation cohort was subsequently applied to the adjusted APC scores in the main cohort (*n*=64) for subgroup classification.

### Human MRI Scanner Hardware and sMRI Data Acquisition

Structural MRI (sMRI) data acquisition took place at the Cerebral Imaging Center of the Douglas Mental Health University Institute Research Centre (Montréal, QC, Canada) between 2012-2023. For all participants, baseline sMRI brain images from 2012 to 2017 were performed on a Siemens MAGNETOM 3T Tim Trio system. Starting in 2018, sMRI data were collected on an upgraded version of the same system (Siemens MAGNETOM 3T Prisma Fit). Both hardware (i.e., gradient, radiofrequency (RF), shimming) and software components of the system were included in this upgrade. In the present study, human sMRI data were used for intermodal imaging coregistration and custom population-template building.

For sMRI data collected on the Siemens MAGNETOM 3T Tim Trio system, T1-weighted (T1w) images were acquired with a 3D magnetization prepared rapid gradient echo (MP-RAGE) sequence with the following parameters: matrix dimensions = 256 x 240 x 176; voxel size = 1 mm x 1 mm x 1 mm; repetition time (TR) = 2300 ms; echo time (TE) = 2.98 ms; inversion time (TI) = 900 ms; flip angle (FA) = 9°; phase encode = anterior-posterior (A>>P); Bandwidth (BW) = 240 Hertz/pixel; GeneRalized Autocalibrating Partially Parallel Acquisitions (GRAPPA) 2; acquisition time (TA) = 5.12 min. For sMRI data collected on the upgraded system (Siemens MAGNETOM 3T Prisma Fit), T1w images were acquired with a 3D MP-RAGE sequence encoding similar parameters: matrix dimensions = 256 x 256 x 192; voxel size = 1 mm x 1 mm x 1 mm; TR = 2300 ms; TE = 2.96 ms; TI = 900 ms; FA = 9°; phase encode = A>>P; BW = 240 Hertz/pixel; slice thickness = 1 mm; GRAPPA 2; TA = 5.30 min.

### Longitudinal sMRI Data Processing

Preprocessing of human sMRI data was performed using the SPM12 toolbox (https://www.fil.ion.ucl.ac.uk/spm/software/spm12/) in MATLAB 2021b. A two-step method for longitudinal registration was implemented using the *serial longitudinal pipeline* from the SPM12 toolbox^106^, as described previously^101,107^. First, the number of T1w scans acquired for each participant was determined. In the case of multiple T1w images within a single time point, the serial longitudinal pipeline was used to create an *intra-scan* average of the corresponding images. Next, an *inter-scan* midpoint average image from the collection of single or averaged T1w images was computed for each participant. The midpoint average T1w images were then segmented with the Computational Anatomy Toolbox’s (CAT12)^108^ *segmentation* module using tissue priors enhanced to improve the classification of subcortical grey matter^109^. The imported grey and white matter segments from the midpoint average T1w images were used to create a custom, age-appropriate cohort template by implementing diffeomorphic registration using geodesic shooting^110^. A deformation field between the cohort-specific template and each participant’s midpoint average T1w image was also estimated during this step and then applied without modulation to the midpoint average T1w images to warp them into the cohort-specific template space. To ensure the integrity of all midpoint average T1w images for intermodal coregistration, a weighted-average Image Quality Rating (IQR) score was computed in CAT12^108^. Images with a CAT12 IQR below 70% were flagged for manual inspection. Finally, the cohort-specific template was nonlinearly registered to the Human Connectome Project Multi-Modal Parcellation-extended (HCP MMPex) atlas space^111^.

### Human PET Imaging Timeline

Cross-sectional VAChT-PET ([^18^F]-FEOBV) data were collected in main cohort participants from 2020-2023. All main cohort participants (*n*=64) and a subset of validation cohort participants (*n*=173) also underwent tau-PET ([^18^F]-AV1451) and Aβ-PET ([^18^F]-NAV4694) imaging between 2017-2023. Descriptive information for PET interscan intervals in the main and validation cohorts are provided in **Table 1**. Subgroup comparisons of the interscan intervals for both cohorts are presented in **Table 2**. Participant-level timelines of VAChT-PET, tau-PET, and Aβ-PET data collection in the main cohort are shown in **Extended Data Fig. 1**.

### Human PET Radiotracer Synthesis

Radiotracer synthesis for human PET imaging took place on each scan date in the McConnell Brain Imaging Center’s (BIC) Cyclotron Facility at McGill University (Montréal, QC, Canada). [^18^F]-FEOBV was synthesized using a pure *levo* precursor (ABX advanced biochemical compounds, Radberg, DE) to obtain the (−)-FEOBV enantiomer, which is the only stereochemical configuration that exhibits binding affinity to VAChT^112^. ^18^F labeling of (−)-FEOBV was performed in accordance with a previous study at the same site^20^. [^18^F]-NAV4694 (Navidea Biopharmaceuticals, Dublin, OH) and [^18^F]-AV1451 (Eli Lilly & Co, Indianapolis, IN) were synthesized and labelled in the same facility as previously described^113^.

### Human PET Scanner Hardware and Data Acquisition

All PET imaging took place in the BIC PET unit (Montréal, QC, Canada) on an ECAT High Resolution Research Tomograph (HRRT, CTI/Siemens) PET scanner. Prior to each participant’s PET scans, a time-standardized transmission scan was acquired. For all participants, [^18^F]-FEOBV brain PET imaging was conducted across two 30-minute imaging sessions, performed with a timing window of 6 ns and a 400–650 keV discrimination energy range^114^. The first session began following the intravenous injection of [^18^F]-FEOBV and continued for 30 minutes (0–30 minutes; data not used in this study). Participants were then removed from the scanner before undergoing a second imaging session. This second session consisted of a 30-minute PET scan (180–210 minutes post-injection; 6 frames; 5 minutes frame ^-1^), corresponding to the established time period for [^18^F]-FEOBV data acquisition in AD patients^20^. [^18^F]-NAV4694 and [^18^F]-AV1451 PET scans were acquired 40–70 minutes (6 frames; 5 minutes frame ^-1^) and 80–100 minutes (4 frames; 5 minutes frame ^-1^) post-injection, respectively. All PET images were reconstructed using an iterative, 3D ordered subset expectation maximization (OSEM3D) algorithm with 10 iterations and 16 subsets and included corrections for 3D scattering^115^. Each frame was reconstructed to a final dimension of 256 x 256 x 207 and a voxel size of 1.22 mm x 1.22 mm x 1.22 mm.

### Human PET Data Processing

Human VAChT ([^18^F]-FEOBV), tau ([^18^F]-AV1451) and Aβ ([^18^F]-NAV) brain PET data were processed using the SPM12 toolbox (https://www.fil.ion.ucl.ac.uk/spm/software/spm12/) in MATLAB 2021b, as described previously^101,107^. Briefly, images were visually inspected to ensure proper orientation and centered to the anterior commissure. For each participant, an *intra-tracer* average of frames was computed, and the average images were coregistered to the corresponding native space midpoint average T1w image. The coregistered average PET images were then normalized to each participant’s mean uptake within previously validated radiotracer-dependent reference regions (VAChT: supratentorial-supraventricular white matter^20,116^; Aβ: cerebellar grey matter^113^; tau: inferior cerebellar grey matter^113^) to produce standardized uptake value ratio (SUVr) images. The deformation fields created during the custom sMRI template building were used to warp the SUVr images without modulation into the cohort-specific template space.

### Normative Deviation Mapping

To characterize participant-specific variations in PET radiotracer uptake, a normative deviation technique was adapted from prior work^45,46^. This method involves two key steps: (1) the computation of population median and deviation reference images and (2) the segmentation of individual voxel-wise data into discrete regions of interest (ROI).

#### Reference Image Computation

To compute reference images, the preprocessed PET data for each participant was taken as input. Given the age-range and unequal proportion of males and females in the main cohort (**Table 1**), a general linear model with age and sex as covariates was used to adjust each individual’s preprocessed PET images for these sources of variation. Additional preprocessing steps performed for normative deviation mapping included masking the PET data by a BFpz ROI. The BFpz ROI was created by combining the HCP MMPex^111^ atlas parcels for all cortical, hippocampal, amygdalar, and BF regions into a binary mask, non-linearly warping this mask to the respective population template space, and taking the logical overlap of this mask with the population template grey matter segmentation. The masked PET data were then smoothed using a 2 mm full-width at half-maximum (FWHM) Gaussian kernel.

For each tracer, reference images were separately generated in a leave-one-out cross-validation framework. First, the SUVr PET image of the target participant was excluded, and the remaining participants’ SUVr PET images were stacked to form a 4D dataset (i.e., participants × voxels). Voxel-wise summary statistics were then computed across the 4D dataset, yielding a reference median (*M-i*) image and a median absolute deviation (MAD) image (*MAD-i*) for each iteration. For normally distributed data, the MAD systematically underestimates the standard deviation. To correct for this, a scaling factor (*S*) of ∼1.4826 was applied to the MAD, making it a consistent estimator of the standard deviation under the assumption of normality^117^. All *M*-*i* and *MAD*-*i* reference images were stored as 3D-volumetric images.

#### Deviation-Based Voxel Segmentation

Using the reference images (*M*-*i* and scaled *MAD*-*i*), each participant’s PET data were segmented into four non-overlapping ROIs based on voxel-wise deviations from the reference images. The median deviation ROIs are: (1) below the lower 95% CI (<L95), (2) between the lower 95% CI and median (L95–M), (3) between the median and upper 95% CI (M–U95), and (4) above the upper 95% CI (>U95). For each segmentation step, continuous and binarized voxel masks were generated. Continuous maps retained the original voxel intensities, while binarized maps indicated voxel inclusion/exclusion. All SUVr extractions from normative deviation ROIs were performed using the binarized maps on the unsmoothed, adjusted PET images.

### Reference Image Stability Analysis

To assess the stability of the population reference images (*M*-*i* or *MAD*-*i*) and independence from individuals characterized by strong deviations from the rest of the population, correlations were performed for each pair of the computed *M-i* or *MAD-i* reference images. This led to a distribution of Pearson *r* correlation coefficients based on (*n* x *n*-1)/2 = 2016 comparisons, given *n*=64 individuals. Descriptive statistics for these analyses are reported in **Supplemental Table 1**.

### Voxel-Wise Regressions of Human Triple PET Radiotracer Dataset

To perform within-participant voxel-wise regressions, SUVr PET images for VAChT, tau, and Aβ were first masked using the binary BFpz ROI. Voxel values within the mask were then extracted, converted to vectors, and independently *z*-scored for each radiotracer, resulting in a normalized distribution of radiotracer uptake values for each participant. Linear regressions were conducted using the voxel vectors for each individual:

1. *VAChT_v_* = *β*0_*indiv*_ + *β*1_*indiv*_ · *Tau*_*v*_ + *β*2_*indiv*_ · *Aβ*_*v*_ + *ε*_*v*_
2. *Aβ*_*v*_ = *β*0_*indiv*_ + *β*1_*indiv*_ · *Tau*_*v*_ + *β*2_*indiv*_ · *VAChT*_*v*_ + *ε*_*v*_

where *V* is the voxel-wise PET image, *β*0 is the intercept, and *ε* is the unexplained variance.

### Adaptive and Lag-Loss Spatial Overlap Models

To identify regions where VAChT is disproportionately high or low relative to tau, Jaccard similarity values were computed between each individual’s VAChT and tau normative deviation mapping ROIs (<L95, <L95-M, >M-U95, >U95; referred to as ROI_1_-ROI_4_ below). These values were then used to defined two biologically motivated models of spatial mismatch. In the *adaptive* model, VAChT was shifted one deviation level above tau, consistent with a cholinergic presynaptic response that exceeds local tau burden. In the *lag-loss* model, tau was shifted one deviation level above VAChT, consistent with a relative failure of cholinergic presynaptic adaptation. Operationally, the adaptive model comprises the ROI pairs tau-ROI_1_ with VAChT-ROI_2_, tau-ROI_2_ with VAChT-ROI_3_, and tau-ROI_3_ with VAChT-ROI_4_, whereas the lag-loss model comprises VAChT-ROI_1_ with tau-ROI_2_, VAChT-ROI_2_ with tau-ROI_3_, and VAChT-ROI_3_ with tau-ROI_4_. Importantly, the voxels contributing to the adaptive and lag-loss models are spatially independent. For each individual, Jaccard similarity values from the constituent ROI pairs were averaged to yield a summary statistic for each model. To localize where the overlaps of each model were most highly expressed in the brain, the Jaccard similarity values were computed at the voxel-level for each participant. The overlap of the within-participant Jaccard similarity values were then calculated for each parcel of the BFpz from the HCP MMPex atlas^111^. Finally, the parcel overlap values were compared between resilient and vulnerable subgroups using the Wilcoxon rank-sum test, adjusting for the FDR (< 0.05) across parcels.

### Aβ-PET and Tau-PET Positivity Thresholds and Classification

To provide clinical context on the PREVENT-AD cohort participants, the percentage of individuals previously identified^118^ as Aβ-PET or tau-PET positive are reported in **Table 1**. Briefly, participants were classified as Aβ-PET positive if their SUVr in a global Aβ index ROI was ≥1.265. Tau-PET positivity was defined as a tau SUVr in a meta-temporal ROI equal to or exceeding the mean tau SUVr of Aβ-PET negative participants plus two standard deviations, corresponding to a threshold of 1.291. Visualization of the ROIs used for Aβ-PET and tau-PET positivity calculations can be found in **Supplemental Fig. 3**. Subgroup comparisons of these measures are presented in **Table 2**.

### Plasma Biomarker Data

Plasma samples collected in 2021-2023 (**Table 1,2**) were available for all main cohort PREVENT-AD participants (*n*=64) and a subset of the validation cohort (*n*=224). All plasma biomarkers were measured with evaluators blinded to clinical information at the Centre for Studies in the Prevention of Alzheimer’s disease (McGill University, Montréal, QC, Canada) on the Simoa SR-X platform (Quanterix, Billerica, MA, USA). Preanalytical preparation for plasma was described previously^119^ with commercially available AD biomarkers kits. Plasma Aβ_1-40_, Aβ_1-42_, and total tau were measured using the Neurology 3-plex A Advantage assay kit, whereas phosphorylated tau at threonine-181 (ptau_181_) was assessed using the new pTau-181 Advantage V2.1 kit from Quanterix (Billerica, MA, USA). All measurements were performed as per manufacturer instructions using internal and external standards.

### Mouse Ethics Statement

All small-animal imaging procedures were conducted in accordance with the Canadian Council of Animal Care’s current policies and were approved by the University of Western Ontario’s Animal Care Committee (Animal Use Protocols: 2020-163) and the Lawson Health Research Institute’s Health and Safety board. Mice were housed as pairs in standard plexiglass cages in temperature- and humidity-controlled rooms (22–25°C, 40%–60%) on 12-hour light/12-hour dark cycles, where food (regular chow) and water were made available *ad libitum*. Imaging and touchscreen experiments were performed during the light cycle (7:00 AM–7:00 PM). All efforts were made to maximize animal welfare before, during, and after experiments.

### Mouse Multimodal Imaging Experimental Overview

High-resolution *in vivo* 9.4T sMRI and [^18^F]-FEOBV microPET data were acquired at the Center of Functional and Metabolic Mapping (Agilent Animal MRI Scanner, Bruker) and the Lawson Health Research Institute’s Preclinical Imaging Facility (Inveon DPET, Siemens Healthineers), respectively. All imaging data were acquired in mice at 6–7 months of age. Data were acquired in *wt* mice (*wt*: MRI = 8 male, 6 female; PET = 5 male, 6 female), either C57BL/6 J or VAChT^flox/flox^, VAChT^Nk2x.1-Cre-flox/flox^ mice (forebrain VAChT *ko*: MRI = 5 male, 5 female; PET = 4 male, 3 female), Nkx2.1-Cre mice (*Cre*-only: MRI = 3 male, 3 female), and ChAT-ChR2-eYFP mice (*hyp*: MRI/PET = 4 male, 3 female). An independent analysis of the *wt* [^18^F]-FEOBV microPET dataset (*n*=11) was performed in the reporting laboratories’ previously published study^120^. VAChT^flox/flox^ and VAChT^Nk2x.1-Cre-flox/flox^ mice were generated on a C57BL/6 J background as previously described^28,61,121^. Nkx2.1-Cre (C57BL/6J-Tg(Nkx2-1-cre)2Sand/J) and ChAT-ChR2-eYFP mice (B6.Cg-Tg(Chat-COP4*H134R/EYFP)6Gfng/J) were purchased from The Jackson Laboratory (JAX: stock no. 008661, stock no. 014546), with the latter maintained as hemizygous.

### Mouse sMRI Data Acquisition

Mice were anesthetized with 1.5–2.0% isoflurane in 1.5 L min^-1^ O_2_ (g) and then moved to the center of a 30 mm RF volume coil (Bruker). Four iterations of 3D magnetization transfer (MT)-weighted spoiled gradient-recalled echo (GRE) images were acquired using a fast low-angle shot pulse sequence with the following parameters: FOV = 16 mm x 16 mm x 10 mm; matrix dimensions = 128 x 128 x 8; voxel size = 0.125 mm x 0.125 mm x 0.125 mm, slice thickness = 10 μm; TR = 30 ms; TE = 2.77 ms; FA = 9°; off-resonance saturation = 4.5 kHz (Gaussian pulse = 10 ms duration); effective FA = 900°^122^.

### Mouse sMRI Data Processing

Preprocessing of mouse anatomical brain images was performed using the SPM12 toolbox (https://www.fil.ion.ucl.ac.uk/spm/software/spm12/) in MATLAB 2020b as described previously^120^. Anatomical images (*n*=4) were averaged, resized, centered, and re-oriented to follow the right-hand coordinate system (RAS). Images were then coregistered to the mouse brain template from Hikishima *et al.*^123^, bias-corrected using MICO^124^, and segmented using tissue priors for grey matter, white matter, and cerebrospinal fluid tissue compartments^123^. The grey and white matter segmentations were then used to create nonlinear transformations for diffeomorphic registration to a study-specific average template using geodesic shooting^110^. Finally, the deformation fields estimated during the template-building step were applied with modulation to each mouse’s grey matter segment to warp the images to template space. To parcellate mouse anatomical (and [^18^F]-FEOBV, see below) brain images, the study-specific mouse brain template was non-linearly registered to the Allen Mouse Brain Atlas (ABA) common coordinate framework version 3 (ABAccfv3)^125^. For *wt*, forebrain VAChT *ko*, and *Cre*-only mice, brain regions for grey matter volume extractions were defined according to the symmetrized, coarse annotated ABAccfv3^125^. Extractions were restricted to voxels within each mouse’s grey matter tissue segment that had a probability of at least 0.1 of belonging to the grey matter. Grey matter volume extractions were performed on the unsmoothed, modulated-warped grey matter segments, adjusted for each mouse’s total intracranial volume (TIV), which was calculated by summing the grey matter, white matter and cerebrospinal fluid volumes in the *seg8.mat file created during segmentation^126^.

### Mouse [^18^F]-FEOBV microPET Data Acquisition

*In vivo,* dynamic [^18^F]-FEOBV microPET data were acquired in *wt*, forebrain VAChT *ko*, and *hyp* mice, as previously described^120^. All [^18^F]-FEOBV microPET imaging sessions were performed as 150-minute list-mode emission scans, with a timing window of 3.432 ns and a 350–640 keV discrimination energy range. Mice were anesthetized with 1.5–2.0% isoflurane in 1.0 L min^-1^ O_2_ (g) and then injected intravenously with ∼20 Megabecquerel (MBq) of [^18^F]-FEOBV. Data were binned into 15 frames of 40 seconds, followed by 28 frames of 300 seconds. An OSEM3D algorithm with two iterations and 18 subsets was used to reconstruct [^18^F]-FEOBV microPET images with the following parameters: matrix dimensions: 128 × 128 × 159; voxel size: 0.776 mm × 0.776 mm × 0.796 mm.

### Mouse [^18^F]-FEOBV microPET Data Processing

Parametric total volume of distribution (V_T_) images for each mouse’s [^18^F]-FEOBV data were estimated as described previously^120^. Whole-brain [^18^F]-FEOBV time-activity curves were fit according to Logan *et al*.^127^, where an image-derived input function (IDIF) from the lumen of the left ventricle of the heart served as the reference^120^. Preprocessing of mouse [^18^F]-FEOBV V_T_ images was performed using the SPM12 toolbox (https://www.fil.ion.ucl.ac.uk/spm/software/spm12/) in MATLAB 2020b. Preparation of [^18^F]-FEOBV V_T_ images included resizing, centering, and re-orienting the images to follow the RAS. Each V_T_ image was then coregistered and cropped to the bounding box of its corresponding native space MRI image, and warped without modulation to the study-specific anatomical mouse brain template. For *wt* and *hyp* mice, extractions of average [^18^F]-FEOBV uptake in the mouse BFpz, hippocampus and cortex were performed using region-specific masks based on the ABAccfv3^125^. A similar approach was also taken to create masks for extracting average [^18^F]-FEOBV uptake in the forebrain (BFpz and striatum) and brainstem (midbrain, pons, and medulla) of *wt* and forebrain VAChT *ko* mice. In all cases, average uptake was calculated on the unsmoothed, warped [^18^F]-FEOBV V_T_ brain images.

### Immunoblot Analysis

Brain tissue samples for Western blotting were collected in a subset of *wt*, forebrain VAChT *ko*, and *hyp* mice following [^18^F]-FEOBV microPET scans. Mice were anesthetized via intraperitoneal injection of a ketamine (100 mg kg^-1^) - xylazine (25 mg kg^-1^) 0.9% sodium chloride solution and then euthanized by cervical dislocation. Brain tissue was extracted, rapidly dissected on ice into appropriate tissue compartments (i.e., brainstem, striatum, hippocampus, cortex), snap-frozen on liquid nitrogen, and stored in 2 ml microcentrifuge tubes at -80°C until needed. Protein isolation and immunoblotting were performed as previously described^128^. Briefly, tissue samples were thawed on ice and homogenized in RIPA buffer (50 mM Tris, 150 mM NaCl, 0.1% SDS, 0.5% sodium deoxycholate, 1% triton X-100, pH 8.0), supplemented with phosphatase inhibitors (1 mM NaF and 0.1 mM Na_3_VO_4_) and a protease inhibitor cocktail (diluted 1:100, Cat#539134-1SET; Calbiochem). Homogenates were sonicated, rocked for 20 minutes at 4°C, and then centrifuged at 10,000g for 20 minutes at 4°C. Supernatants were collected and protein quantification was performed using the Pierce™ BCA Protein Assay Kit (Cat#23227; ThermoFisher Scientific). 20–25 µg of protein were loaded and separated on 4–12% Bis-Tris Protein Gels (Cat#NW04125BOX; Invitrogen) and then transferred onto PVDF membranes (Cat#IPVH00010; EMD Millipore) using the Trans-Blot Turbo system (Bio-Rad Laboratories, Hercules, CA, USA). Membranes were blocked for 60 minutes in 5% milk (for VAChT) or 5% BSA (for Synaptophysin and β-Actin) in 1xTBST. Primary antibodies included rabbit anti-VAChT (diluted 1:2000, Cat#139103; Synaptic Systems), mouse anti-Synaptophysin (diluted 1:2000, Cat#ab8049; Abcam), and anti-β-Actin peroxidase (diluted 1:25000, Cat#a3854; Sigma-Aldrich). Following 4°C overnight primary antibody incubation in the respective blocking solutions, membranes were washed and then incubated for 60 minutes at room temperature in corresponding horseradish peroxidase (HRP)-conjugated secondary antibodies. Bands were visualized using enhanced chemiluminescence (ECL) detection reagents in a ChemiDoc Imaging System (Bio-Rad Laboratories, Hercules, CA, USA). Densitometric analysis was performed in ImageLab (Bio-Rad Laboratories, Hercules, CA, USA).

### PVD–PVR Touchscreen Testing

7 month-old *wt* (*n*=6) and forebrain VAChT *ko* (*n*=19) mice (male) were tested on the *pvd*–*pvr* task (SOP2; www.touchscreencognition.org) developed for the rodent automated Bussey-Saksida Touchscreen System (Model 81426, Campden Instruments, Lafayette, Indiana) with Abet II Touch Software Version 2.20 (Lafayette Instrument Company, Lafayette, Indiana). The *pvd*–*pvr* task is designed to measure the effects of pharmacological or genetic manipulation on visual learning and cognitive flexibility^129^. Prior to *pvd*–*pvr* testing, all mice completed behavioural habituation, pre-training stages and acquisition sessions in the touchscreen chamber^130^. Following meeting acquisition criteria, mice were subject to *pvd*. The *pvd* sessions require mice to learn to associate a food reward (7 µL strawberry milkshake) with a nose-poke response to a correct image (S^+^ stimulus) in one touchscreen window, while simultaneously ignoring a second visually-distinct, incorrect image (S^−^ stimulus) in the other touchscreen window. After completing six *pvd* sessions (two pre-pump implantation, four post-pump implantation), mice were tested on *pvr* learning across 10 sessions. In *pvr*, the strawberry milkshake food reward becomes linked to the previously incorrect image (S^−^ stimulus), while nose poke responses to the former correct image (S^+^ stimulus) are no longer reinforced.

### Subcutaneous Osmotic Pump Implantation and Pharmacological Manipulation Paradigm

During *pvd*–*pvr* testing, a subset of forebrain VAChT *ko* mice underwent pharmacological manipulation by the continuous administration of the acetylcholinesterase inhibitor, donepezil, via subcutaneous osmotic pumps. Donepezil hydrochloride (C6821-50mg) monohydrate was obtained from Sigma-Aldrich (Oakville, Ontario) and rehydrated using saline and 10% dimethyl sulfoxide (DMSO) to a dose of 1.5 mg kg^-1^. Dosage was determined using the Alzet drug concentration calculator to account for pump model and corresponding flow rate as well as the weight of the mice. Implantation of subcutaneous osmotic pumps (model 1004, Alzet, Durect Corporation, Cupertino, California) was performed in *wt* and forebrain VAChT *ko* mice following completion of PVD session two. Pumps were designed to deliver continuous treatment (*wt* (*n*=6) or forebrain VAChT *ko* (*n*=8): saline + 10% DMSO; forebrain VAChT *ko* (*n*=11): 1.5 mg kg^-1^ of donepezil) at a fixed rate of 0.11µl hour^-1^ for 28 days. Preparation for pump insertion included loading each pump with a 1 ml syringe, capping the pumps with a pin, and soaking them in saline with <1 hour to implantation. Prior to pump insertion, mice were anesthetized with isoflurane (4%, initial; 1.5%, maintenance) in 1 L min^-1^ O_2_ (g) and put in a recumbent prone position. A small incision was made in the upper-right quadrant of each mouse’s back, followed by the insertion of a pump and sealing of the incision site with a wound clip. Mice were given two days to recover from the surgeries before beginning session three of *pvd*.

### Spatial Brain Phenotype Analysis

Regional *t* statistics representing grey matter volume differences between forebrain VAChT *ko* and *wt* mice were computed for 212 anatomical parcels from the ABA, based on the parcellation scheme from Oh *et al*.^65^. Parcels in the brainstem (i.e., midbrain, pons, medulla) and low-intensity olfactory bulb areas were excluded, yielding 144 parcels for analysis. Corresponding regional *in situ* hybridization (ISH) data were obtained from the ABA^66^, preprocessed as described previously^67^. For each parcel, gene expression was represented as "energy" scores, reflecting expression levels weighted by hybridization intensity. To perform module based analyses, a gene-network enriched for presynaptic functions vulnerable to dysregulation in AD was obtained from Supplemental Table 5 in Roussarie *et al*.^58^ (‘module C’, *n*=702). Genes from this network that lacked expression data across an entire parcel were removed. After merging and cleaning, 401 of the 702 genes in the original module remained. A principal-component analysis (PCA) was then applied to the cleaned gene-by-region matrix, and the first principal component was saved. The resulting eigengene score captured the dominant axis of co-expression within the plasticity gene module. Spearman rank correlation coefficients were computed between the region-wise profile of grey matter volume differences and the selected transcriptional profile (i.e., the module eigengene). To assess statistical significance while accounting for spatial autocorrelation, the observed correlation was compared to a null distribution generated from 40,000 spatial surrogate maps^131,132^. These surrogate maps preserved the intrinsic spatial autocorrelation structure of gene expression but randomized regional assignments^67,132^. A two-tailed empirical *P* value was computed as the proportion of null correlations with an absolute value greater than or equal to the observed correlation.

### Whole-Brain snRNAseq Analysis

#### Single-nucleus RNA-sequencing Atlases

Whole-brain adult human (∼3 million nuclei) and mouse (∼6 million nuclei) snRNAseq atlases were downloaded from Siletti *et al*.^56^ and Langlieb *et al*.^57^, respectively. Metadata accompanying each dataset were used to annotate clusters according to cell type (i.e., neuron, glia) and developmental compartment (i.e., telencephalon, diencephalon, mesencephalon, rhombencephalon). Only clusters annotated as neuronal and telencephalic were retained for downstream analyses (human: *n*=235 clusters; mouse: *n*=66 clusters). Raw count matrices were normalized by scaling each nuclei to a total of 10^6^ counts (counts per million, CPM), followed by log_1_p transformation.

#### Dimensionality Reduction

To visualize the anatomical and neurochemical spatial embeddings of telencephalic neuronal populations, highly variable genes (HVGs) were identified from a cluster-size informed subset of 300,000 nuclei in both the adult human and mouse snRNAseq atlases, from which the top 3,000 genes were selected for downstream dimension reduction. HVG selection was performed by computing gene-wise mean and variance across nuclei and ranking genes by dispersion. Dimension reduction was performed using PCA followed by construction of a *k*-nearest neighbour graph and uniform manifold approximation and projection (UMAP). UMAP embeddings were generated from the principal component space using *Scanpy*^133^ and used for visualization and annotation of telencephalic neuronal clusters across both atlases. Neuronal clusters were further annotated according to their primary anatomical origin within the telencephalic BFpz, including the BF, cortex, hippocampus, and amygdala, and other regions, as well as their dominant neurochemical identity (i.e., glutamatergic, GABAergic, cholinergic, other).

#### Gene Module Scoring and Cluster-Level Permutation Testing

Expression of the *a priori p*lasticity gene-network from Roussarie *et al*.^58^ (‘module C’, *n*=702 genes) was first summarized at the single-nuclei level by computing the mean normalized expression of genes within the module^134,135^. Cluster-level enrichment of the plasticity gene-network module was then evaluated across all telencephalic neuronal clusters^136,137^. When cross-referencing the plasticity gene-network list with the snRNAseq datasets, 671 (95.6%, human) and 650 (92.6%, mouse) genes were found to overlap. Cluster-level module scores were obtained by aggregating nuclei-level scores within each telencephalic neuronal cluster using the median expression across nuclei. Enrichment of the plasticity gene-network module in each telencephalic neuronal cluster was then assessed by comparing the cluster-level module score to the pooled module score across all other telencephalic neurons in the atlas (Δ=cluster score-pooled score). Statistical significance was determined using permutation testing in which cluster labels were randomly shuffled across nuclei while preserving cluster sizes to generate empirical null distributions of module scores. For each permutation, cluster-level module scores were recomputed and compared with pooled scores to generate a null distribution of enrichment statistics. This procedure was repeated 1,000 times to estimate empirical *P* values for each cluster. Adjustment for the False Discovery Rate (FDR < 0.05) was then applied using the Benjamini-Hochberg (BH) procedure.

#### Gene-Level Enrichment Analysis

Gene-level analyses were performed to identify individual genes enriched in BF cholinergic neurons (human: SPLAT_400; mouse: MC_12) relative to other telencephalic neuronal populations. For each gene in the transcriptome, mean expression in the target cholinergic cluster was compared with mean expression across all remaining telencephalic neurons. Permutation testing was used to assess statistical significance. For each permutation, a set of nuclei equal in size to the cholinergic target cluster was randomly sampled from the atlas to generate a null target population, and the difference (Δ) in mean expression between the sampled target nuclei and the remaining nuclei was computed. This procedure was performed across 59,480 genes in the human dataset and 21,899 genes in the mouse dataset. Empirical *P* values were calculated by comparing the observed expression difference to the permutation-derived null distribution, and multiple testing correction was performed using the BH procedure across empirical *P* values for all genes to obtain transcriptome-wide FDR adjusted estimates.

#### Over-Representation Analyses

Confirmatory over-representation analysis of plasticity network genes overlapping with the human and mouse snRNAseq datasets was performed using *SynGO* (v1.3), an evidence-based, expert-curated resource of synapse-specific Gene Ontology (GO) terms^59^. The plasticity network genes intersecting the human or mouse snRNAseq datasets were used as the foreground gene lists. The full set of genes in the human or mouse snRNAseq datasets were used as the background lists. For both datasets, 16 GO biological processes (BP) terms were identified as significantly enriched (FDR < 0.01), with a minimum of 10 matching input genes.

Exploratory over-representation analysis was performed to investigate whether specific transcriptional programs within the plasticity gene-network were enriched in BF cholinergic neuros. Here, the plasticity network genes exhibiting significant positive enrichment (Δ > 0, *q* < 0.05 FDR) in humans or mice relative to all other neuronal populations were used as the foreground gene lists. The complete set of plasticity network genes intersecting the human or mouse snRNAseq datasets were used as the background gene lists. The over-representation analysis was conducted using *ShinyGO* (v0.85.1)^138^ to identify enriched GO BPs and cellular component (CC) terms. Enrichment was assessed at FDR < 0.05.

### Statistical Analysis

Statistical analyses were conducted using GraphPad Prism (version 10.6.1; GraphPad Software, La Jolla, CA), except for robust linear regressions and imaging analyses which were performed in MATLAB 2020b or 2021b, and spatial brain phenotyping and snRNAseq analyses, which were carried out in Python v3.13.2. Sample sizes were determined based on previous studies in the field. Parametric tests and their corresponding post hoc comparisons were applied to groups with a sample size of at least 10 that met required visual and/or statistical assumptions. When these assumptions were not met, non-parametric tests with appropriate post hoc analyses were used. Statistical test results are provided in the figure legends and **Supplementary Information** files.

## Acknowledgments

This work was supported by the Canadian Institutes for Health Research (193293, 453677) to H.R.C.S. and T.W.S., the Alzheimer’s Society of Canada Research Program (grant 176677, project ID 22-11) to T.W.S. and K.M.O., the Healthy Brains for Healthy Lives initiative, Alzheimer’s Association (AARG-22-927100) and a NIA R01 (AG068563) to R.N.S. Funding for PREVENT-AD was initiated through a $13.5 million, seven-year public-private partnership, in 2011. Support was received from McGill University, the Fonds de recherche du Québec-Santé, an unrestricted research grant from Pfizer Canada, the J.L. Levesque Foundation, the Douglas Hospital Research Centre and its Foundation, the Government of Canada, and the Canada Foundation for Innovation. Private sector contributions were facilitated by the Development Office of McGill University’s Faculty of Medicine and the Douglas Hospital Research Centre Foundation. The PREVENT-AD cohort is continued to be supported in part by grants from the Canadian Institutes for Health Research (J.P., S.V.), the Fonds de recherche du Québec-Santé (R.N.S., J.P., S.V.) and the J.L. Levesque Foundation. T.W.S. holds the Cecil and Linda Rorabeck Chair in Molecular Neuroscience and Vascular Biology (Western University, London, ON, Canada). M.A.M.P. is a Tier 1 Canada Research Chair in Neurochemistry (Western University, London, ON, Canada). The authors would like to thank Miranda Bellyou, Dr. Alex Li, and Joe Gati from the CFMM, Lise Desjardins and Jennifer Hadway from Lawson Imaging, as well as Matthew Cowan, Sanda Raulic and Jue Fan for their involvements in the small animal imaging experiments and their animal breeding and animal husbandry efforts, respectively. We would also like to thank Dr. Leon French (University of Toronto, Toronto, ON, Canada) for his feedback on the manuscript. Finally, we extend our gratitude to the PREVENT-AD study participants and their families for their commitment and dedication to the research program, as well as the staff and core members of the PREVENT-AD research group. The list of PREVENT-AD members can be found at https://preventad.loris.ca/acknowledgements/acknowledgements.php?date=%5b2020-08-05.

## Contributions

V.F.P., M.A.B., M.A.M.P., R.N.S., and T.W.S obtained the funding for the study. K.M.O., M.A.M.P., R.N.S., and T.W.S. conceived the project, designed the study, and interpreted the results. K.M.O., and T.W.S. ran and generated all main analyses and figures. M.A.M.P., R.N.S., and T.W.S. supervised the project. Preclinical data collection and experimentation was performed by L.A.D., Q.Q., A.M.C., L.G-C., F.H.B., M.S.F., J.W.H., and K.M.O. Pre-processing and analysis of the preclinical data was performed by L.A.D., Q.Q., L.G-C., and K.M.O. K.M.W., T.Q., A.F-V., J.R., A.W., G.R.T., E.A., J.T-M., J-P.S., J.P., and K.M.O. contributed to the collection, organization, and curation of the human datasets. T.Q., A.F-V., H.R.C.S., R.A.M.H., K.M.O., and T.W.S. pre-processed and analyzed the human imaging dataset. G.G.N., T.S.A., K.M.O., and T.W.S. contributed to the organization and analysis of the open-source transcriptomic datasets. K.M.O., R.N.S., and T.W.S. wrote the initial draft of the manuscript with contributions from H.R.C.S., R.A.M.H., T.S.A., J.D.T., M.S.F., J.W.H., T.J.B., J-P.S., J.P., V.F.P., M.A.M.P., and S.V.P. All authors contributed their scientific expertise to the final manuscript.

## Data and Materials Availability

Human demographic, cognition, and imaging data are from the PREVENT-AD program, which is promoting data access for the scientific community via their Canadian Alzheimer Platform (Brain Canada Foundation). For more information regarding PREVENT-AD data usage and availability, please refer to https://www.centre-stopad.com/en/plateforme-cap. The ABA ISH data^66^ is publicly available for download (https://community.brain-map.org/t/in-situ-hybridization-ish-data/2762). The human^56^ (https://alleninstitute.github.io/abc_atlas_access/descriptions/WHB_dataset.html) and mouse^57^ (https://docs.braincelldata.org/downloads) snRNAseq atlases are publicly available for download. The list of genes used to compute plasticity gene-network scores (‘module C’) can be found in Table S5 from Roussarie *et al*.^58^. All [^18^F]-FEOBV microPET datasets will be made available on Zenodo for download upon publication of the manuscript. Mouse anatomical images are under active use by the reporting laboratories and can be made available by contacting. All code will be made available upon publication. Source data for all figures are in the corresponding **Source Data** files.

## Extended Data

**Extended Data Fig. 1.**
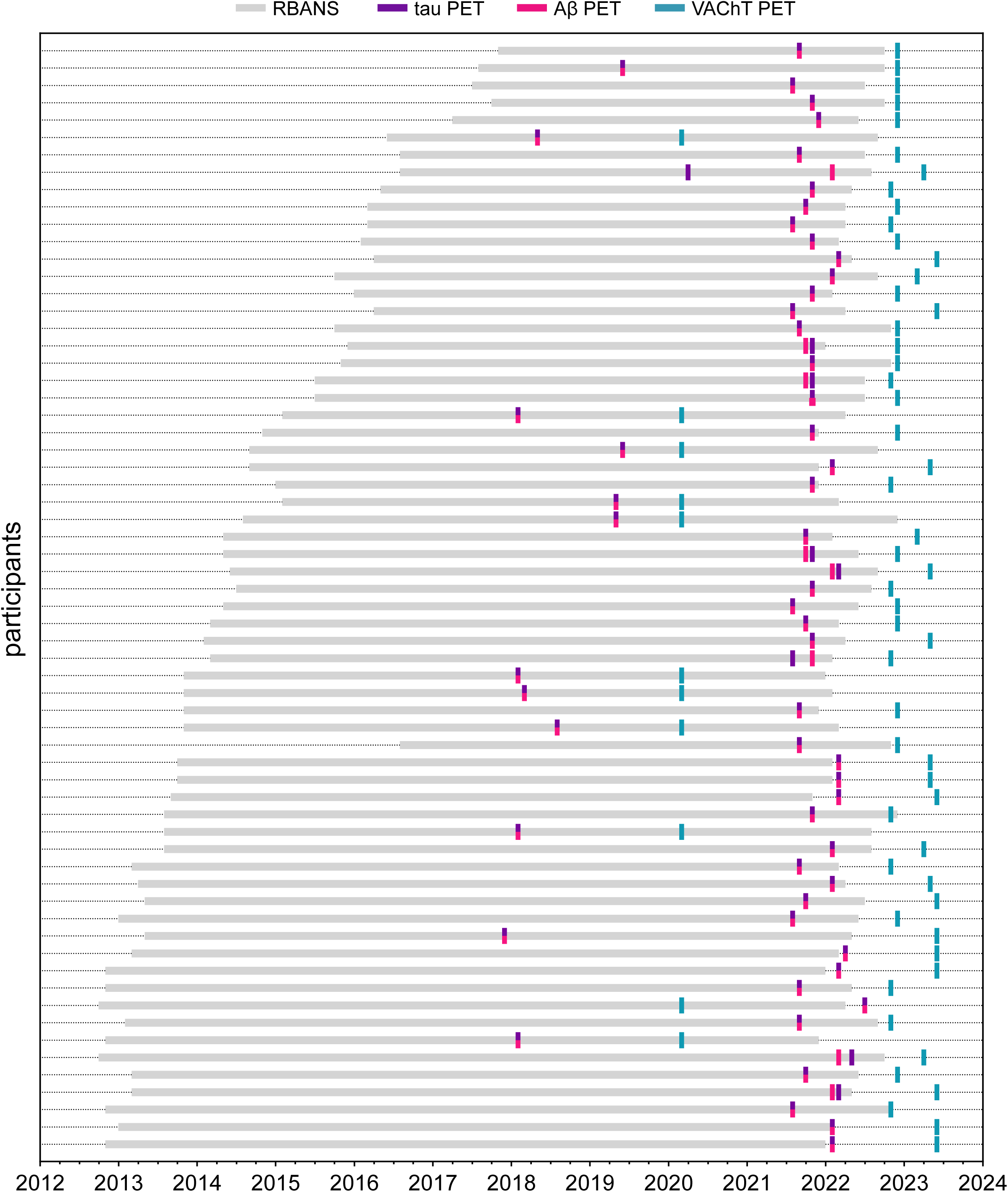
PREVENT-AD main cohort PET and RBANS individual data collection timelines. Participant-level timelines of PET imaging and RBANS evaluations in main cohort (*n*=64) PREVENT-AD participants. Vertical tick marks indicate the timing of VAChT-PET (teal), tau-PET (purple), and Aβ-PET (pink) scans. The horizontal grey bars represent the interval between baseline and final RBANS evaluations. Timelines are displayed in months.

**Extended Data Fig. 2.**
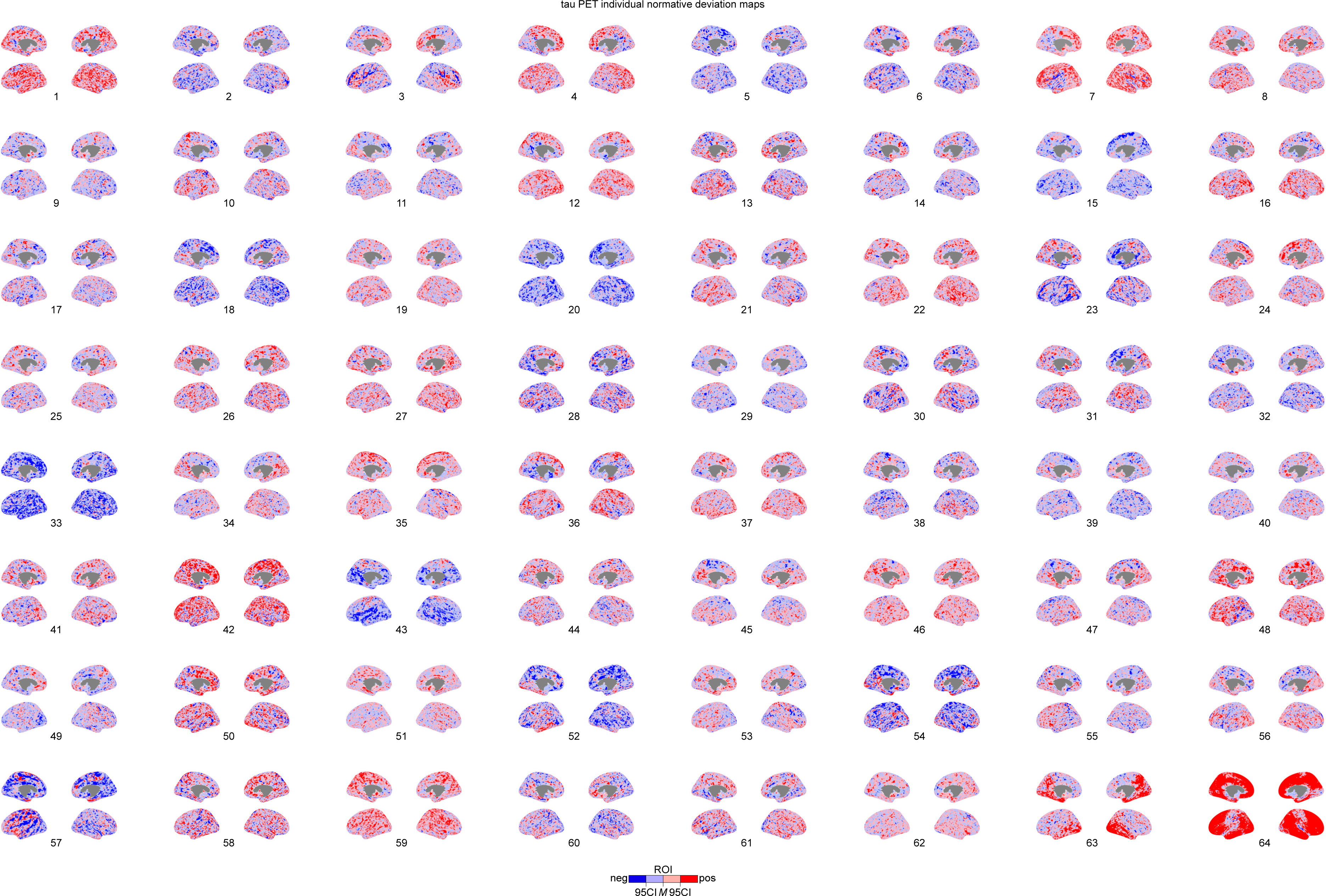
Individual tau-PET normative median deviation mapping profiles. Segmented normative deviation regions of interest (ROI) for tau-PET images *(n*=64), projected on medial and lateral brain surfaces. ROIs are colour coded according to the corresponding confidence interval (CI) bin (<L95=dark blue; L95–M=light blue; M–U95=light red; >U95=dark red).

**Extended Data Fig. 3.**
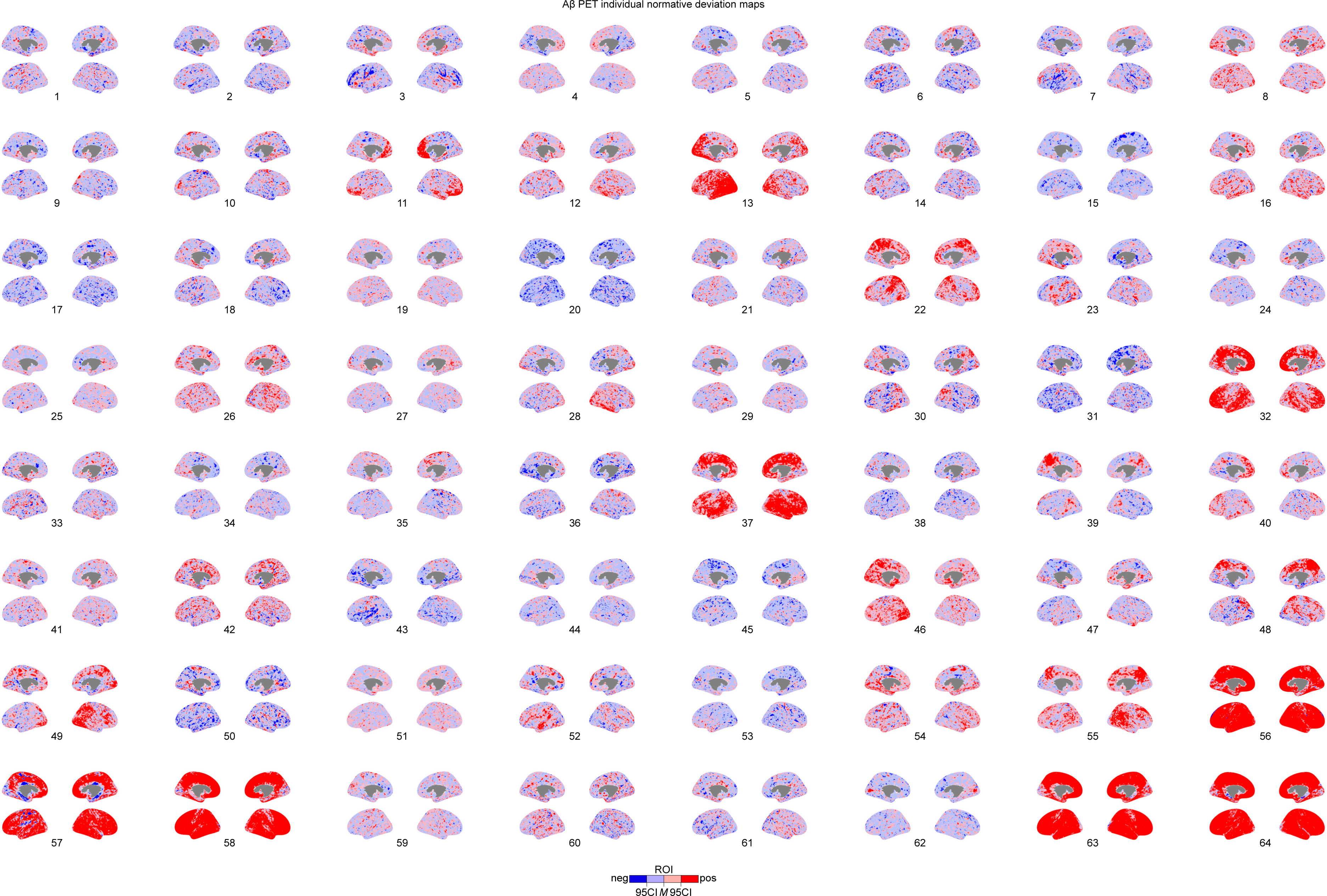
Individual Aβ-PET normative median deviation mapping profiles. Segmented normative deviation regions of interest (ROI) for Aβ-PET images *(n*=64), projected on medial and lateral brain surfaces. ROIs are colour coded according to the corresponding confidence interval (CI) bin (<L95=dark blue; L95-M=light blue; M-U95=light red; >U95=dark red).

**Extended Data Fig. 4.**
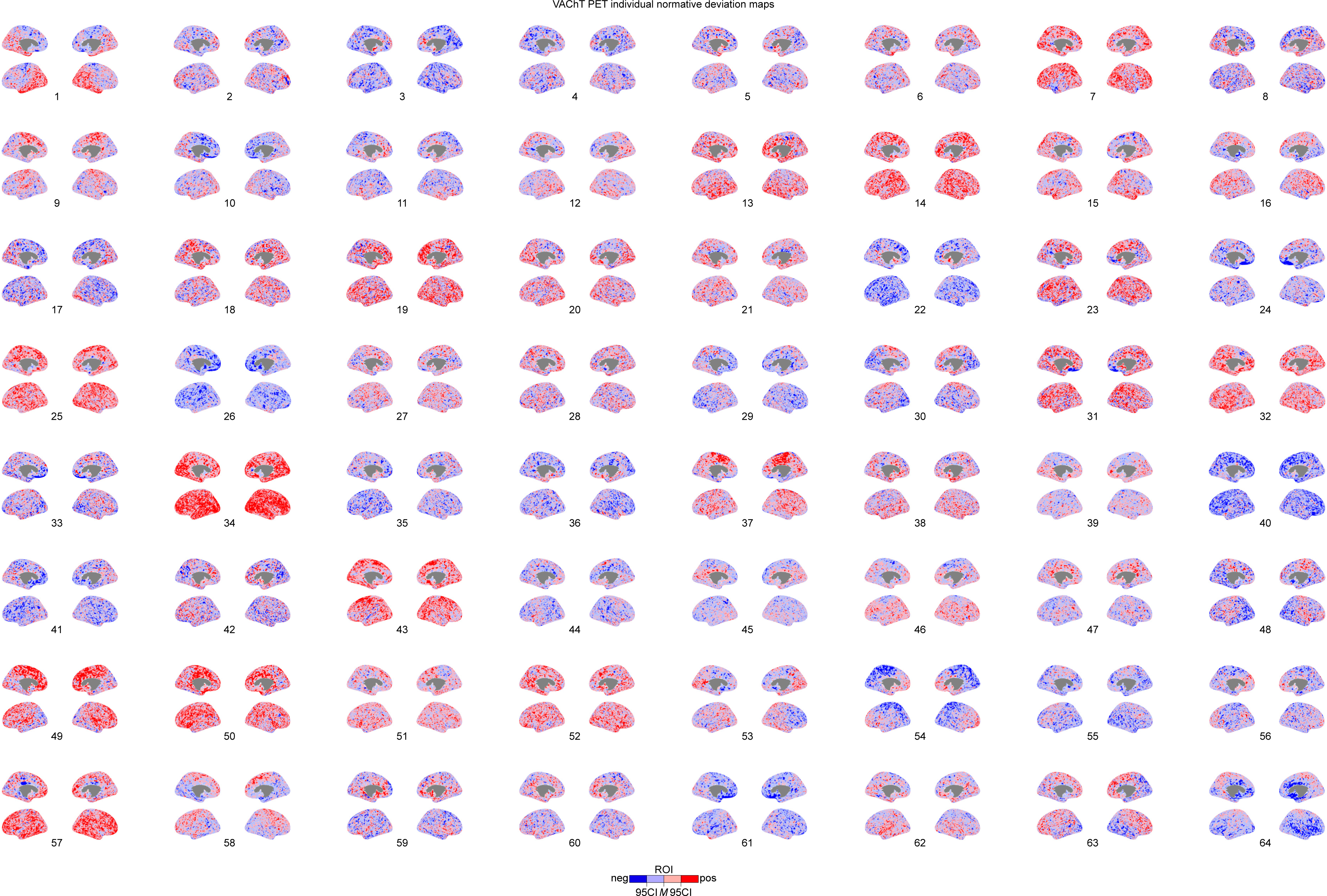
Individual VAChT-PET normative median deviation mapping profiles. Segmented normative deviation regions of interest (ROI) for VAChT-PET images *(n*=64), projected on medial and lateral brain surfaces. ROIs are colour coded according to the corresponding confidence interval (CI) bin (<L95=dark blue; L95-M=light blue; M-U95=light red; >U95=dark red).

**Extended Data Fig. 5.**
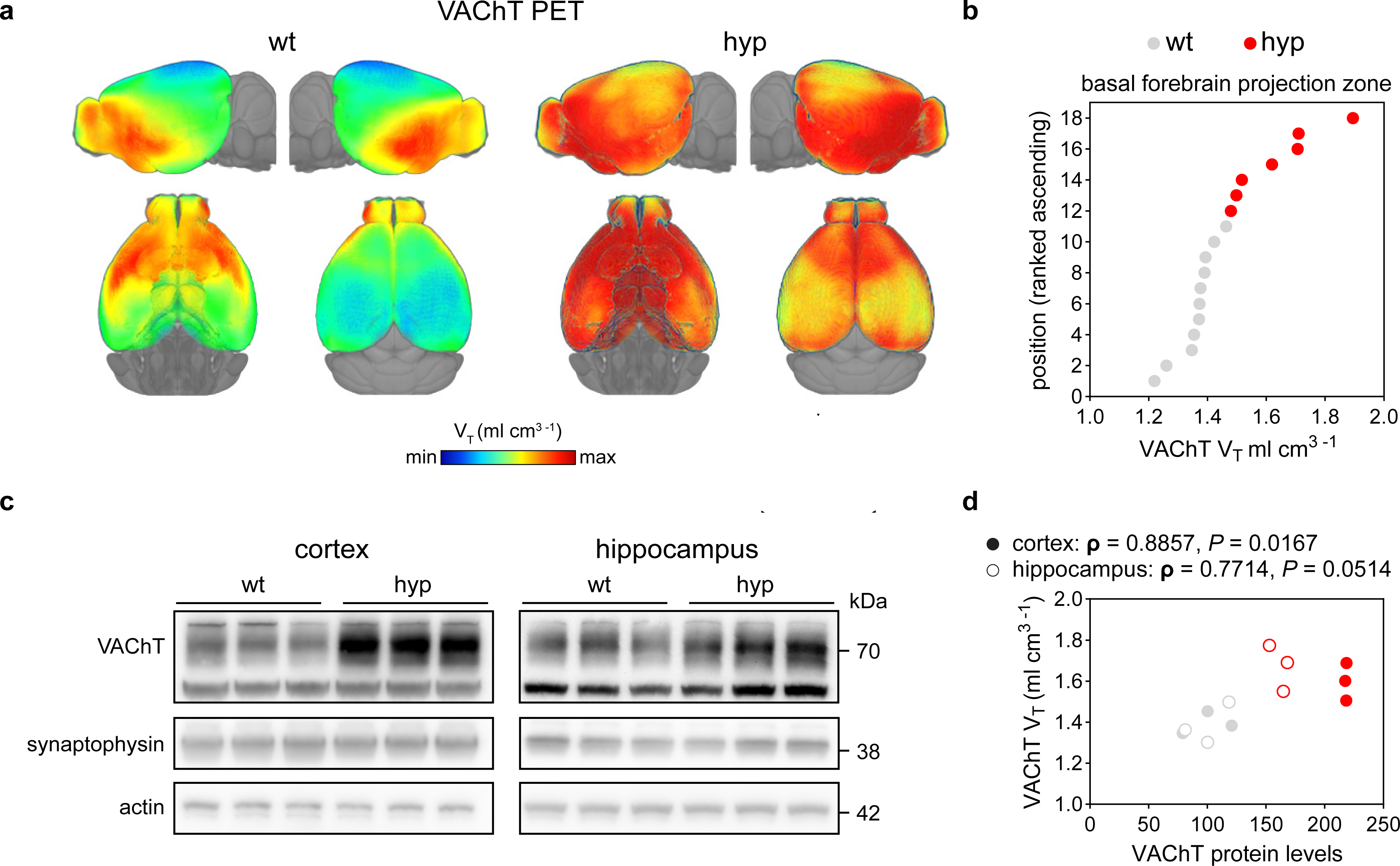
[^18^F]-FEOBV uptake reflects endogenous increases in synaptic VAChT levels. **a**, Representative [^18^F]-FEOBV total volume of distribution (V_T_) images for *wt* (*left*) and *hyp* (*right*) mice (min-max=0.3–2.1). **b**, Scatter plot showing individual mean [^18^F]-FEOBV V_T_ values in the mouse BFpz, ranked from lowest to highest (*wt*: *n*=11, grey; *hyp*: *n*=7, red). **c**, Immunoblot analysis of VAChT protein levels in *wt* (*n*=3) and *hyp* (*n*=3) mice in the cortex (*left*) and hippocampus (*right*). Synaptophysin and actin were used as protein loading controls. **d**, Scatter plot between VAChT protein levels from **c** in cortical (closed circle) and hippocampal (open circle) tissue in a subset of *wt* and *hyp* mice, and [^18^F]-FEOBV V_T_ values in the corresponding brain region (one-sided Spearman rank correlations). Example mouse data displayed using MRIcroGL: https://www.nitrc.org/projects/mricrogl. Full membranes for **c** are displayed in **Supplemental Fig. 1**. Source data in **Source Data** file.

**Extended Data Fig. 6.**
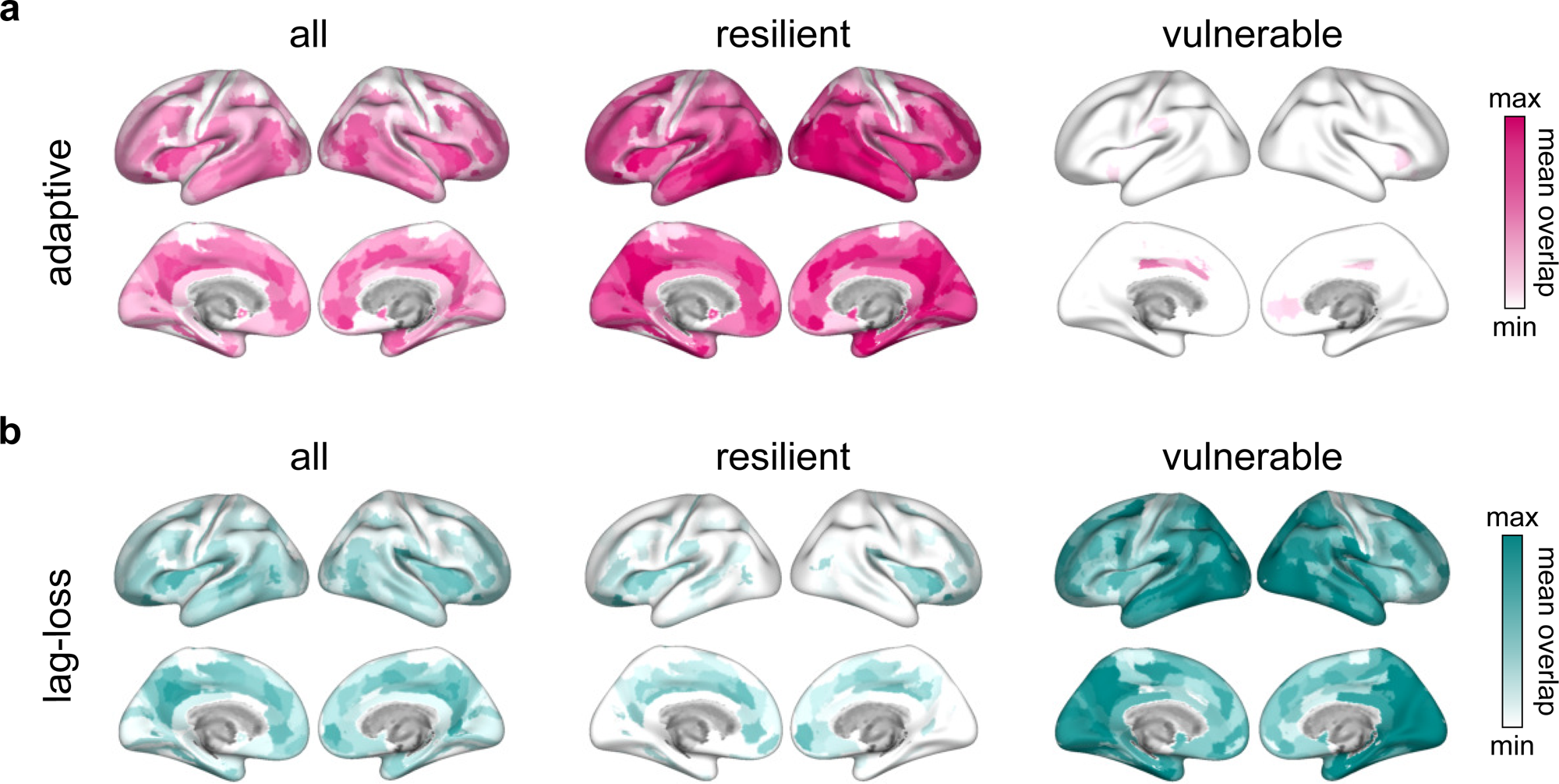
Adaptive and lag-loss models of tau-VAChT colocalization in healthy older adults at risk for Alzheimer’s disease. **a,b**, Mean overlap of voxel-level Jaccard similarity values with parcels in the BFpz for adaptive (**a**) and lag-loss (**b**) models across all participants (*left*, *n=*64; min-max=0.2–0.25), resilient individuals (*middle*, *n*=52; min-max=0.2–0.25), and vulnerable individuals (*right*, *n*=12; min-max=0.2–0.3), displayed on medial and lateral brain surfaces. Parcels are from the Human Connectome Project Multi-Modal Parcellation-extended atlas^111^. Source data in **Source Data** file.

