## Supplementary Information File for "Cholinergic synaptic plasticity shapes resilience and vulnerability to tau"

**Supplemental Tables**

Supplemental Table 1. Descriptive Statistics for Reference Image Stability Analysis.

Supplemental Table 2. Statistical information for Figure 1b.

Supplemental Table 3. Statistical information for Figure 1f.

Supplemental Table 4. Statistical information for Figure 1g.

Supplemental Table 5. Statistical information for Figure 1h.

Supplemental Table 6. Statistical Information for Figure 2b.

Supplemental Table 7. Statistical Information for Figure 2c.

Supplemental Table 8. Statistical Information for Figure 2p (in .xlsx file).

Supplemental Table 9. Statistical Information for Figure 2q (in .xlsx file).

Supplemental Table 10. Statistical Information for Figure 3d (in .xlsx file).

Supplemental Table 11. Statistical Information for Figure 3g (in .xlsx file).

Supplemental Table 12. Statistical Information for Figure 4a.

Supplemental Table 13. Statistical Information for Figure 4b.

Supplemental Table 14. Statistical Information for Figure 4c.

Supplemental Table 15. Statistical Information for Figure 4d.

Supplemental Table 16. Statistical Information for Figure 4h.

Supplemental Table 17. Statistical Information for Figure 4i (in .xlsx file).

Supplemental Table 18. Statistical Information for Figure 4i Supplementary Analysis (in .xlsx file).

Supplemental Table 19. Statistical Information for Figure 4j (in .xlsx file).

**Supplemental Figures**

Supplemental Figure 1. Full Membranes of Western Blots in Extended Data Figure 5c.

Supplemental Figure 2. Full Membranes of Western Blots in Figure 4a-d.

Supplemental Figure 3. Meta ROIs for Calculation of Aβ-PET and tau-PET Positivity.

**Supplemental Table 1. Descriptive Statistics for Reference Image Stability Analysis.** Descriptive statistics (median, range) for stability analysis of tau-PET, Aβ-PET, and VAChT-PET normative deviation mapping reference images. Median and range are given for *n*=2016 comparisons per PET reference image.

| PET radiotracer | reference image | median *r* | range (minimum–maximum) |
|---|---|---|---|
| **tau** | M-i | 0.99953 | 0.99946–0.99974 |
| | MAD-i | 0.98998 | 0.98868–0.99167 |
| **Aβ** | M-i | 0.99959 | 0.99949–0.99997 |
| | MAD-i | 0.99033 | 0.98739–0.99862 |
| **VAChT** | M-i | 0.99982 | 0.99976–0.99991 |
| | MAD-i | 0.99488 | 0.99401–0.99549 |

**Supplemental Table 2. Statistical information for Figure 1b.** Results of one sample Wilcoxon signed-rank test on Jaccard similarity values between like normative deviation mapping ROIs (<L95, L95-M, M-U95, >U95) for tau-PET and Aβ-PET data.

| percent Jaccard similarity (tau, Aβ) | | | | |
|---|---|---|---|---|
| **one sample Wilcoxon signed-rank test** | **<L95** | **L95-M** | **M-U95** | **>U95** |
| **theoretical median** | 0 | 0 | 0 | 0 |
| **actual median** | 3.245 | 31.97 | 26.82 | 2.105 |
| **number of values** | 64 | 64 | 64 | 64 |
| **sum of signed ranks (*W*)** | 2080 | 2080 | 2080 | 2016 |
| ***P* value (two-tailed)** | <0.0001 | <0.0001 | <0.0001 | <0.0001 |

**Supplemental Table 3. Statistical information for Figure 1f.** Results of Friedman test with Dunn’s multiple comparisons test comparing (*top*) number of voxels between tau¬Aβ ROIs (<L95 vs. L95-M; M-U95 vs. >U95) as well as (*middle*) tau and (*bottom*) Aβ SUVr levels across tau¬Aβ ROIs (<L95 → L95-M → M-U95 → >U95).

| **tau¬Aβ: number of voxels** | | | |
|---|---|---|---|
| **Friedman test** | **Friedman statistic** | | ***P* value (approximate)** |
| | 149.8 | | <0.0001 |
| **Dunn's multiple comparisons test** | **Δrank** | ***Z*** | ***P* value (two-tailed)** |
| **L95-M - <L95** | 130 | 8.900 | <0.0001 |
| **M-U95 - >U95** | 120 | 8.216 | <0.0001 |

| **tau¬Aβ: tau SUVr** | | | |
|---|---|---|---|
| **Friedman test** | **Friedman statistic** | | ***P* value (approximate)** |
| | 192 | | <0.0001 |
| **Dunn's multiple comparisons test** | **Δrank** | ***Z*** | ***P* value (two-tailed)** |
| **L95-M - <L95** | 64 | 4.382 | <0.0001 |
| **M-U95 - L95-M** | 64 | 4.382 | <0.0001 |
| **>U95 - M-U95** | 64 | 4.382 | <0.0001 |

| **tau¬Aβ: Aβ SUVr** | | | |
|---|---|---|---|
| **Friedman test** | **Friedman statistic** | | ***P* value (approximate)** |
| | 114 | | <0.0001 |
| **Dunn's multiple comparisons test** | **Δrank** | ***Z*** | ***P* value (two-tailed)** |
| **L95-M - <L95** | 140 | 9.585 | <0.0001 |
| **M-U95 - L95-M** | -125 | 8.558 | <0.0001 |
| **>U95 - M-U95** | 58 | 3.971 | 0.0002 |

**Supplemental Table 4. Statistical information for Figure 1g.** Results of Friedman test with Dunn’s multiple comparisons test comparing (*top*) number of voxels between Aβ¬tau ROIs (<L95 vs. L95-M; M-U95 vs. >U95) as well as (*middle*) tau and (*bottom*) Aβ SUVr levels across Aβ¬tau ROIs (<L95 → L95-M → M-U95 → >U95).

| **Aβ¬tau: number of voxels** | | | |
|---|---|---|---|
| **Friedman test** | **Friedman statistic** | | ***P* value (approximate)** |
| | 131.1 | | <0.0001 |
| **Dunn's multiple comparisons test** | **Δrank** | ***Z*** | ***P* value (two-tailed)** |
| **L95-M - <L95** | 139 | 9.517 | <0.0001 |
| **M-U95 - >U95** | 89 | 6.093 | <0.0001 |

| **Aβ¬tau: tau SUVr** | | | |
|---|---|---|---|
| **Friedman test** | **Friedman statistic** | | ***P* value (approximate)** |
| | 141.5 | | <0.0001 |
| **Dunn's multiple comparisons test** | **Δrank** | ***Z*** | ***P* value (two-tailed)** |
| **L95-M - <L95** | 141 | 9.654 | <0.0001 |
| **M-U95 - L95-M** | -151 | 10.34 | <0.0001 |
| **>U95 - M-U95** | 87 | 5.956 | <0.0001 |

| **Aβ¬tau: Aβ SUVr** | | | |
|---|---|---|---|
| **Friedman test** | **Friedman statistic** | | ***P* value (approximate)** |
| | 192 | | <0.0001 |
| **Dunn's multiple comparisons test** | **Δrank** | ***Z*** | ***P* value (two-tailed)** |
| **L95-M - <L95** | 64 | 4.382 | <0.0001 |
| **M-U95 - L95-M** | 64 | 4.382 | <0.0001 |
| **>U95 - M-U95** | 64 | 4.382 | <0.0001 |

**Supplemental Table 5. Statistical information for Figure 1h.** Results of Friedman test with Dunn's multiple comparisons test comparing (*top*) number of voxels between tau∩Aβ ROIs (<L95 vs. L95-M; M-U95 vs. >U95) as well as (*middle*) tau and (*bottom*) Aβ SUVr levels across tau∩Aβ ROIs (<L95 → L95-M → M-U95 → >U95).

| **tau∩Aβ: number of voxels** | | | |
|---|---|---|---|
| **Friedman test** | **Friedman statistic** | | ***P* value (approximate)** |
| | 142.1 | | <0.0001 |
| **Dunn's multiple comparisons test** | **Δrank** | ***Z*** | ***P* value (two-tailed)** |
| **L95-M - <L95** | 138 | 9.448 | <0.0001 |
| **M-U95 - >U95** | 106 | 7.257 | <0.0001 |

| **tau∩Aβ: tau SUVr** | | | |
|---|---|---|---|
| **Friedman test** | **Friedman statistic** | | ***P* value (approximate)** |
| | 189 | | <0.0001 |
| **Dunn's multiple comparisons test** | **Δrank** | ***Z*** | ***P* value (two-tailed)** |
| **L95-M - <L95** | 63 | 4.347 | <0.0001 |
| **M-U95 - L95-M** | 63 | 4.347 | <0.0001 |
| **>U95 - M-U95** | 63 | 4.347 | <0.0001 |

| **tau∩Aβ: Aβ SUVr** | | | |
|---|---|---|---|
| **Friedman test** | **Friedman statistic** | | ***P* value (approximate)** |
| | 189 | | <0.0001 |
| **Dunn's multiple comparisons test** | **Δrank** | ***Z*** | ***P* value (two-tailed)** |
| **L95-M - <L95** | 63 | 4.347 | <0.0001 |
| **M-U95 - L95-M** | 63 | 4.347 | <0.0001 |
| **>U95 - M-U95** | 63 | 4.347 | <0.0001 |

**Supplemental Table 6. Statistical information for Figure 2b.** Results of linear mixed-effects model (REML) with Fisher's least significant difference test comparing baseline and final RBANS total scores from the resilient and vulnerable subgroups in the PREVENT-AD validation cohort.

| **validation cohort: baseline and final RBANS total score by subgroup** | | |
|---|---|---|
| **mixed-effects model (REML)** | ***F* (DFn, DFd)** | ***P* value** |
| **time** | 195.2 (1, 283) | <0.0001 |
| **group** | 40.94 (1, 283) | <0.0001 |
| **time x group** | 125.0 (1, 283) | <0.0001 |

| **Fisher's LSD difference test** | | **LS Δmean** | **SE** | ***t*** | ***P* value (two-tailed)** |
|---|---|---|---|---|---|
| **vulnerable - resilient** | **baseline** | -1.51 | 1.63 | 0.924 | 0.3561 |
| | **final** | -17.31 | 1.63 | 10.611 | <0.0001 |
| **baseline - final** | **vulnerable** | 17.78 | 1.31 | 13.565 | <0.0001 |
| | **resilient** | 1.97 | 0.53 | 3.723 | 0.0002 |

**Supplemental Table 7. Statistical information for Figure 2c.** Results of linear mixed-effects model (REML) with Fisher's least significant difference test comparing baseline and final RBANS total scores from the resilient and vulnerable subgroups in the PREVENT-AD main cohort.

| **main cohort: baseline and final RBANS total score by subgroup** | | |
|---|---|---|
| **mixed-effects model (REML)** | ***F* (DFn, DFd)** | ***P* value** |
| **time** | 49.1 (1, 62) | <0.0001 |
| **group** | 1.01 (1, 62) | 0.3177 |
| **time x group** | 39.9 (1, 62) | <0.0001 |

| **Fisher's LSD test** | | **LS Δmean** | **SE** | ***t*** | ***P* value (two-tailed)** |
|---|---|---|---|---|---|
| **vulnerable - resilient** | **baseline** | 3.54 | 2.93 | 1.209 | 0.2289 |
| | **final** | -9.10 | 2.93 | 3.103 | 0.0024 |
| **baseline - final** | **vulnerable** | 13.30 | 1.80 | 7.387 | <0.0001 |
| | **resilient** | 0.69 | 0.86 | 0.798 | 0.4277 |

**Supplemental Table 8. Statistical Information for Figure 2p.** Results of Wilcoxon rank-sum test comparing Jaccard similarity values from the adaptive tau-VAChT model across Human Connectome Project extended atlas parcels in resilient versus vulnerable PREVENT-AD main cohort participants. Full statistics are provided in the accompanying .xlsx file.

**Supplemental Table 9. Statistical Information for Figure 2q.** Results of Wilcoxon rank-sum test comparing Jaccard similarity values from the lag-loss tau-VAChT model across Human Connectome Project extended atlas parcels in resilient versus vulnerable PREVENT-AD main cohort participants. Full statistics are provided in the accompanying .xlsx file.

**Supplemental Table 10. Statistical Information for Figure 3d.** Results of permutation testing for cluster-level plasticity gene-network scoring in telencephalic neurons in the healthy adult human brain. Full statistics are provided in the accompanying .xlsx file.

**Supplemental Table 11. Statistical Information for Figure 3g.** Results of permutation testing for cluster-level plasticity gene-network scoring in telencephalic neurons in the healthy adult mouse brain. Full statistics are provided in the accompanying .xlsx file.

**Supplemental Table 12. Statistical information for Figure 4a.** Results of Mann-Whitney *U* test comparing synaptophysin protein levels in cortical samples from wild-type (*wt*) and forebrain VAChT knock-out (*ko*) mice.

| **cortex: synaptophysin protein levels** | | |
|---|---|---|
| **Mann-Whitney *U* test** | ***U*** | ***P* value (two-tailed)** |
| **wt - ko** | 6 | 0.6857 |

**Supplemental Table 13. Statistical information for Figure 4b.** Results of Mann-Whitney *U* test comparing synaptophysin protein levels in hippocampal samples from wild-type (*wt*) and forebrain VAChT knock-out (*ko*) mice.

| **hippocampus: synaptophysin protein levels** | | |
|---|---|---|
| **Mann-Whitney *U* test** | ***U*** | ***P* value (two-tailed)** |
| **wt - ko** | 7 | 0.8857 |

**Supplemental Table 14. Statistical information for Figure 4c.** Results of Mann-Whitney *U* test comparing synaptophysin protein levels in striatal samples from wild-type (*wt*) and forebrain VAChT knock-out (*ko*) mice.

| **striatum: synaptophysin protein levels** | | |
|---|---|---|
| **Mann-Whitney *U* test** | ***U*** | ***P* value (two-tailed)** |
| **wt - ko** | 5 | 0.4857 |

**Supplemental Table 15. Statistical information for Figure 4d.** Results of Mann-Whitney *U* test comparing synaptophysin protein levels in brainstem samples from wild-type (*wt*) and forebrain VAChT knock-out (*ko*) mice.

| **brainstem: synaptophysin protein levels** | | |
|---|---|---|
| **Mann-Whitney *U* test** | ***U*** | ***P* value (two-tailed)** |
| **wt - ko** | 2 | 0.1143 |

**Supplemental Table 16. Statistical information for Figure 4h.** Results of one-sample Wilcoxon signed-rank test for pairwise visual discrimination (pvd) and reversal (pvr) performance in saline-treated wild-type (*wt*) mice, saline-treated forebrain VAChT knock-out (*ko*) mice, and donepezil-treated (1.5 mg $kg^{-1}$) forebrain VAChT *ko* mice.

| **Wilcoxon signed-rank test** | | | **wt (saline)** | | **ko (saline)** | | **ko (donepezil)** | |
|---|---|---|---|---|---|---|---|---|
| **$H_0$: $x \leq 50$** | | | ***W*** | ***P* value** | ***W*** | ***P* value** | ***W*** | ***P* value** |
| **off** | **pvd** | **session 1** | 21 | 0.0156 | 36 | 0.0039 | 66 | 0.0005 |
| | | **session 2** | 21 | 0.0156 | 36 | 0.0039 | 66 | 0.0005 |
| **on** | | **session 3** | 21 | 0.0156 | 36 | 0.0039 | 66 | 0.0005 |
| | | **session 4** | 21 | 0.0156 | 36 | 0.0039 | 66 | 0.0005 |
| | | **session 5** | 21 | 0.0156 | 36 | 0.0039 | 66 | 0.0005 |
| | | **session 6** | 21 | 0.0156 | 36 | 0.0039 | 66 | 0.0005 |
| | **pvr** | **session 7** | 0 | 1.0000 | 1 | 0.9961 | 0 | 1.0000 |
| | | **session 8** | 5 | 0.8906 | 3 | 0.9883 | 0 | 1.0000 |
| | | **session 9** | 4 | 0.9219 | 6 | 0.9609 | 15 | 0.9492 |
| | | **session 10** | 9 | 0.4063 | 1.5 | 0.9844 | 12.5 | 0.9385 |
| | | **session 11** | 17 | 0.0938 | 7 | 0.9492 | 40 | 0.1094 |
| | | **session 12** | 20 | 0.0313 | 0 | 1.0000 | 43.5 | 0.1934 |
| | | **session 13** | 17.5 | 0.0938 | 13.5 | 0.3281 | 35.5 | 0.2158 |
| | | **session 14** | 21 | 0.0156 | 13 | 0.5938 | 55 | 0.0010 |
| | | **session 15** | 21 | 0.0156 | 12.5 | 0.4063 | 45 | 0.0020 |
| | | **session 16** | 21 | 0.0156 | 15 | 0.4609 | 66 | 0.0005 |

**Supplemental Table 17. Statistical information for Figure 4i.** Results of independent samples *t* test comparing grey matter volumes in parcels from the Allen Mouse Brain Atlas common coordinate framework version 3 in wild-type (*wt*) mice and forebrain VAChT knock-out (*ko*) mice. Full statistics are provided in the accompanying .xlsx file.

**Supplemental Table 18. Statistical Information for Figure 4i Supplementary Analysis.** Results of Wilcoxon rank-sum test comparing grey matter volumes in parcels from the Allen Mouse Brain Atlas common coordinate framework version 3 in wild-type (*wt*) mice and Nkx2.1-Cre (*Cre*-only) mice. Full statistics are provided in the accompanying .xlsx file.

**Supplemental Table 19. Statistical information for Figure 4j.** Results of independent samples *t* test comparing grey matter volumes in parcels from the Allen Mouse Brain Atlas in wild-type (*wt*) mice and forebrain VAChT knock-out (*ko*) mice. Full statistics are provided in the accompanying .xlsx file.

**Supplemental Figure 1. Full Membranes of Western Blots in Extended Data Figure 5c.** Uncropped membranes for Western blots of (*top*) VAChT (*middle*) synaptophysin and (*bottom*) actin in the (*left*) cortex and (*right*) hippocampus of wild-type and VAChT hyper-expressing mice.

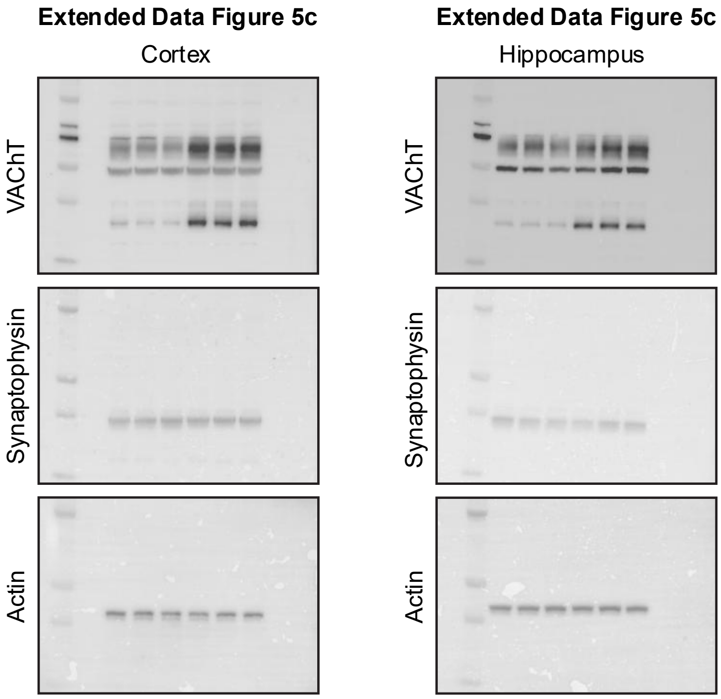

**Supplemental Figure 2. Full Membranes of Western Blots in Figure 4a-d.** Uncropped membranes for Western blots of (*top*) VAChT (*middle*) synaptophysin and (*bottom*) actin in the (*top row*, *left*) cortex, (*top row*, *right*) hippocampus, (*bottom row*, *left*) striatum, (*bottom row*, *right*) brainstem of wild-type and forebrain VAChT knockout mice.

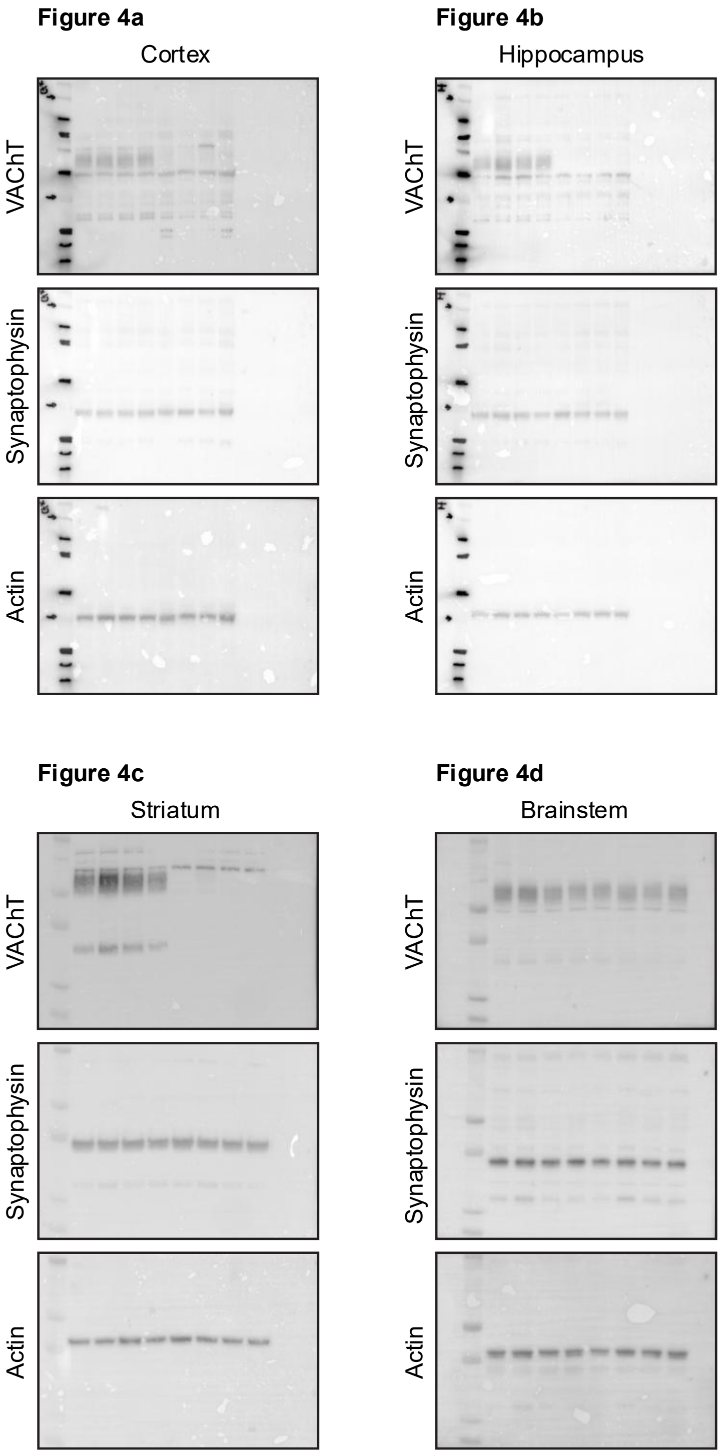

**Supplemental Figure 3. Meta ROIs for Calculation of Aβ-PET and tau-PET Positivity.** Regions of interest (ROI) for calculating (*left*) global Aβ index and (*right*) meta-ROI tau standardized uptake value ratio (SUVr) levels are shown in purple, displayed on medial and lateral surfaces of the brain.

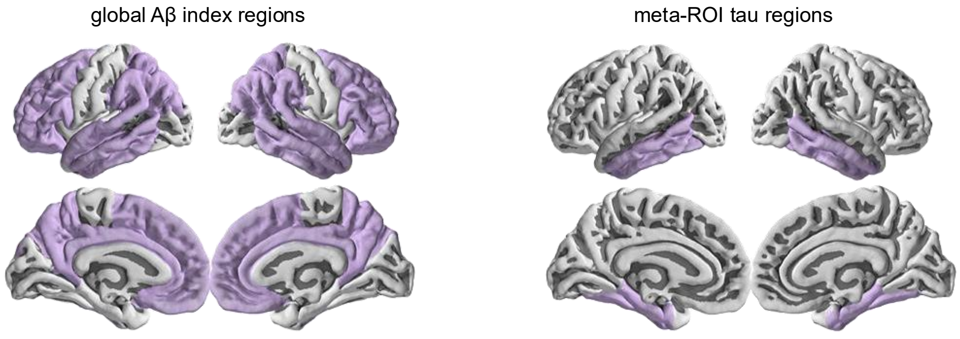
